# POLQ mediates replication-stress induced structural variant formation throughout common fragile sites during mitosis

**DOI:** 10.1101/2024.05.28.596214

**Authors:** Thomas E. Wilson, Samreen Ahmed, Amanda Winningham, Thomas W. Glover

## Abstract

Genomic structural variants (SVs) greatly impact human health and disease, but much is unknown about their generative mechanisms, especially for the large class of nonrecurrent alterations. Common fragile sites (CFSs) are unstable loci that provide a model for SV formation, especially large deletions, under replication stress. We studied SV junction formation as it occurred in cells by applying error-minimized capture sequencing to CFS DNA harvested during replication stress. SV junctions formed throughout CFS genes at a 5-fold higher rate after cells passed from G2 into M-phase. Neither SV formation nor CFS expression depended on mitotic DNA synthesis (MiDAS), an error-prone form of conservative replication active at CFSs. Instead, analysis of tens of thousands of *de novo* SV junctions combined with DNA repair pathway inhibition revealed a primary role for DNA polymerase theta (POLQ)-mediated end-joining (TMEJ) in M-phase SV formation. We propose an important role for TMEJ in nonrecurrent SV formation genome wide.

## Introduction

Chromosomal rearrangements, *i.e.*, structural variants (SVs), represent a large proportion of genomic diversity and are responsible for numerous genomic disorders ^1^. They also arise in somatic cells and are a major mutation type in cancers. A predominant class of SVs are kb- to Mb-scale copy number variants (CNVs), including simple deletions and duplications and more complex rearrangements ^2, 3^. There are two major types of larger (>10kb) CNVs in human genomes as revealed by breakpoint structures. Recurrent CNVs are formed by meiotic unequal recombination between flanking segmental duplications ^4^. Nonrecurrent germline CNVs and virtually all that arise in somatic and cancer cells can form anywhere in the genome and are characterized by short microhomologies or insertions at the breakpoint junctions ^5, 6^.

Despite the large impact of nonrecurrent CNVs on human health, our understanding of the mechanisms responsible for their formation is incomplete. One challenge is that a single mechanism may not explain all SV events. Microhomologies at breakpoint junctions suggested early models of SV formation by nonhomologous end joining (NHEJ) of DNA double-strand breaks (DSBs) ^7^, since microhomologies cannot support the homologous recombination (HR) that drives recurrent CNV formation. However, some non-recurrent CNVs have multiple breakpoint junctions that are difficult to reconcile with NHEJ but might be explained by fork-stalling and template switching (FoSTeS), which invokes a DSB-independent switch of a nascent replication strand to a different template ^8, 9^, or microhomology-mediated break-induced replication (MMBIR), which is similar to FoSTeS but invokes a single-ended DSB at a stalled replication fork ^8, 10^. Importantly, FoSTeS and MMBIR are thought to act during S-phase at sites of stalled replication whereas NHEJ predominates in G1 ^11^.

Over many experiments, we established common fragile sites (CFSs) as an experimental model for non-recurrent CNV formation ^12, 13, 14, 15, 16, 17, 18^. CFSs are genomic loci that show frequent gaps and breaks on metaphase chromosome spreads under replication stress ^18, 19^. CFSs were originally proposed to represent regions of DNA that remained unreplicated past S-phase ^19^ and decades of work supports conclusions that CFSs correspond to the subset of late-replicating genomic loci residing at the largest human genes. When transcribed, these genes become susceptible to impaired replication fork progression due to a paucity of active replication origins resulting from transcription that continues into S-phase ^17, 20, 21^.

CFS loci are also hotspots for CNV formation under replication stress ^17, 22^. Like nonrecurrent CNVs in human genomes, CNVs formed at CFSs are characterized by short microhomologies, insertions, or blunt ends, leading us to propose they could be formed by template switching, NHEJ, or microhomology mediated end joining ^13^. An alternative potential mechanism was suggested by the findings of Minocherhomji et al. ^23^ who showed that unreplicated genome regions complete replication in early mitotic prophase by a rescue process called mitotic DNA synthesis (MiDAS). Notably, peaks of MiDAS synthesis have a high correspondence with CFS genes ^24, 25^. Moreover, MiDAS often results in synthesis on just one sister chromatid suggesting it occurs by conservative break-induced replication (BIR) ^26^, a variant HR pathway operating at single-ended DSBs that is prone to template switching and other errors ^27, 28^. These observations link MiDAS to CFSs, possibly as the mechanism by which SV junctions form at these loci ^29, 30^.

Most recently, a series of studies have described a specific form of end joining catalyzed by DNA polymerase theta (POLQ), a pathway often called theta-mediated end-joining (TMEJ) ^31^. TMEJ creates nonhomologous junctions with a significant frequency of templated insertions that provide microhomologies for bridging two DSB ends. Intriguingly, POLQ activity and TMEJ become activated upon entry into mitosis through mechanisms that include RHINO-directed recruitment to mitotic DNA breaks and POLQ phosphorylation by the PLK1 kinase ^32, 33, 34^. Thus, TMEJ joins NHEJ, FoSTeS, MMBIR, and MiDAS as a candidate mechanism for catalyzing SV junction formation at CFSs and elsewhere.

A limitation of prior literature is a paucity of prospective experimental tests of the hypothesized connections between replication stress, replication rescue and repair mechanisms, and SV formation based on direct assessment of *de novo* SV junctions. The high frequency of CNV formation at CFS loci provides a valuable experimental model for obtaining these important missing breakpoint junction data to move beyond indirect inferences from cancer genomes and cytological studies. We previously used whole genome microarrays to establish that large, actively transcribed CFS genes are hotspots for CNV formation, but throughput and resolution were low and CNVs could only be detected after clonal expansion ^12, 13, 14, 15, 16, 17^. More recently, we established svCapture sequencing as a reliable method for detecting and characterizing locus-specific single-molecule SV junctions ^35^.

Here, we used svCapture to enrich whole-genome sequencing within known CFS genes to determine the distribution, structure, breakpoint junctions, and mechanisms of SV formation during specific cell-cycle phases and in cells deficient in key repair processes. We reasoned that identifying the cell cycle phase at which nonrecurrent SV junctions are formed following replication stress would guide identification of the mechanism(s) involved since FoSTes and MMBIR would likely predominate in S-phase, MiDAS and TMEJ in M-phase, and NHEJ in the following G1 phase. We found that that most replication stress-induced SVs formed at CFSs during M-phase but that MiDAS was not required for their formation. In contrast, by examining SV frequencies in cell populations and the structure of tens of thousands of *de novo* SV junctions, we identified TMEJ as a primary contributor to SV junction formation in M-phase, with NHEJ having a lesser contribution in multiple cell cycle phases. These results reveal a central role for TMEJ in mediating large SV formation and provide important insights into possible genome-wide SV mutation mechanisms in normal and cancer cells.

## Results

### svCapture detects large *de novo* deletion SVs in CFS genes

We used hybridization target capture, *i.e.*, svCapture ^35^, to enrich whole-genome sequencing near the middle of five previously established CFS genes in three cell lines to enable detection of *de novo* single-molecule SV formation in cell populations (**Figure 1A**). Specifically, we targeted large genes *PRKG1*, *NEGR1*, and *MAGI2* in fibroblast line UM-HF1 (HF1) and *FHIT* and *WWOX* in lymphoblastoid line GM12878 and colon cancer line HCT116 as model systems for replication stress-associated SV formation (**Figure S1A** and **S1B**). *PRKG1*, *NEGR1*, and *MAGI2* are known CNV hotspots in HF1 cells ^17^, whereas *FHIT* and *WWOX* are among the loci with the highest frequency of metaphase breaks and gaps in lymphoblastoid and HCT116 cells ^36^.

**Figure 1.**
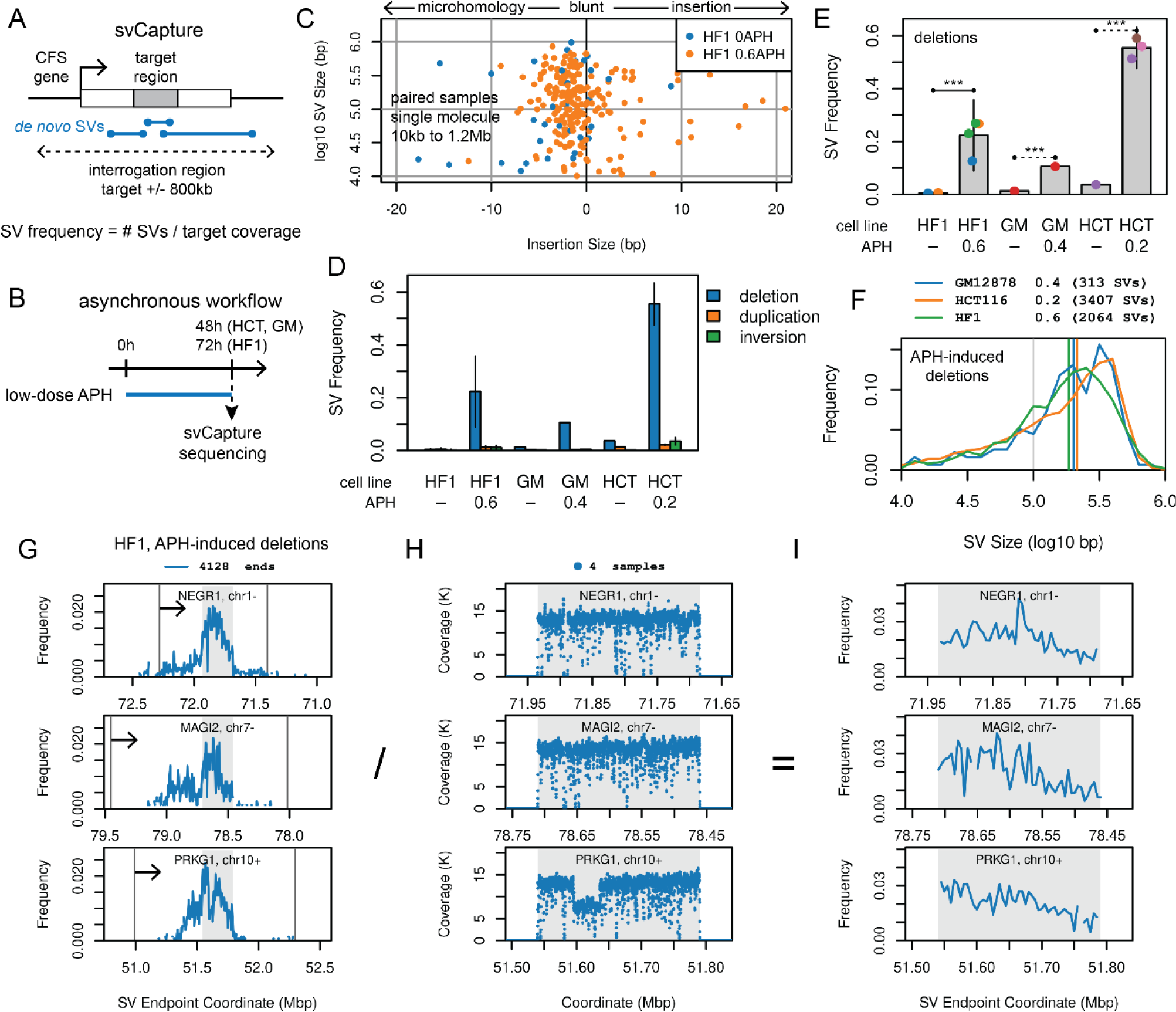
APH-induced SV junctions arise throughout large CFS genes in asynchronous normal and cancer cell lines. **A.** svCapture sequencing was targeted near CFS gene centers to detect SV junctions with at least one breakpoint in a target region. **B.** Timeline of asynchronous cell experiments, with SV induction by low-dose APH. **C.** Example data from paired HF1 samples. Each point is one intrachromosomal, single-molecule SV from 10kb to 1.2Mb. All SVs are plotted in random order; there are many fewer blue points. Negative insertion sizes represent junction microhomologies (see Methods). **D.** Induced SVs are strongly biased to deletions. APH doses are in µM. SV frequency is the junction count divided by the target region fold-coverage. HF1, UM-HF1; GM, GM12878; HCT, HCT116. **E.** Induced deletion SV frequency. Each point aggregates one sample, *e.g.*, all SVs from one color as shown in panel C. Sample point colors denote shared experimental batches. Throughout, all data matching a sample group are plotted, with selected intergroup p-values marked as ns, not significant; *, p <= 0.01; **, p <= 0.001; ***, p <= 0.0001. Dashed significance lines were calculated without over-dispersion due to low sample number. **F.** APH-induced deletion size distributions by cell line. Colored vertical lines are median sizes. **G.** Distribution of APH-induced deletion breakpoints in HF1 cells, which extend beyond the capture targets (shaded) but less than the limits of the interrogation regions (plot width) or gene spans (vertical lines, transcribed left to right, arrow). Data are aggregated in 5kb bins. **H.** High net HF1 target region coverage in 100bp bins, the normalization denominator for panel **I.** Low-coverage bins reflect reduced capture probe density and/or sequencing efficiency. *PRKG1* carried a clonal deletion in the HF1 cells under study at 51.595-51.645Mb. **I.** HF1 deletion breakpoint distributions in target regions after normalizing to read coverage >=500, showing non-focal accumulation of SV junctions.

We first used a workflow in which SVs were allowed to accumulate asynchronously under replication stress caused by low doses of the DNA polymerase inhibitor aphidicolin (APH, **Figure 1B**), determined empirically per cell line to slow but not stop replication and to produce an average of 2-5 chromosome gaps and breaks per cell. Throughout, we only scored SV junctions present in a single source DNA molecule to track *de novo* SV formation during the experiment. Paired samples showed a clear increase in *de novo* SV Frequency, *i.e.*, SV count normalized to target region coverage, in APH-treated vs. control cells (**Figure 1C**). In some cell types, observed frequencies were consistent with as many as half of measured haplotypes acquiring a *de novo* SV (**Figure 1D**), consistent with prior microarray results ^17^.

Induced SVs at CFS genes were strongly biased toward deletions over duplications, inversions, and inter-target translocations (**Figure 1D**), even though svCapture reports all SV junction types detectable by short-read sequencing. The lower level of non-deletion SVs was sometimes significantly induced by APH (**Figure S1C** to **S1G**), but due to the greater abundance of DNA loss originating from unreplicated DNA ^17, 22^, we mainly track deletions below. **Figure 1E** verifies significant and reproducible deletion induction across multiple experimental batches (denoted by point color) in all cell lines.

APH-induced deletion SVs showed mainly short microhomologies at breakpoint junctions and a median SV size of ∼200kb in all cell lines (**Figures 1C**, **1F**, and **S1C**), matching the 186kb median for microarray CNVs in human cell lines ^13, 17^. SVs tended to be smaller without APH induction and for duplications (**Figures S1H** and **S1I**). We therefore applied filters to only track SVs smaller than 1.2Mb and larger than 10kb (50kb for inversions, **Figure S1C**) to maintain specificity.

### SVs formed under replication stress arise throughout large gene bodies

Proposed mechanisms for genomic instability at CFS genes variably invoke replication delay ^17^, local sequence features prone to polymerase stalling within CFS genes ^37, 38^, and additional influences secondary to transcription ^39, 40^. Our high-density collection of SV breakpoints informs this longstanding question. As expected, we saw the greatest frequency of deletion breakpoints in capture target regions (**Figures 1G**, **S2A**, and **S2D**). However, some of the second SV breakpoints were outside the capture targets but remained almost entirely within the gene bodies. Like prior work ^17^, duplications and inversions were more frequently located at the gene flanks as opposed to deletions, which clustered in the center of the genes (**Figure S1D**).

When we normalized deletion breakpoint density within capture targets to local sequencing coverage to account for variable svCapture efficiency and clonal SVs (**Figures 1H**, **S2B**, and **S2E**), we observed that breakpoints were distributed throughout the 250kb or 400kb capture targets (**Figures 1I**, **S2C**, and **S2F**). No specific locations in any of the five targeted genes appeared to act as unexpectedly high frequency sites of focal SV formation. The pattern was consistent with replication forks failing stochastically throughout large, transcribed genes.

Together, svCapture recapitulated all aspects of SV formation at CFS genes under replication stress as previously seen in microarray data but with much greater resolution and data density. We do not know what fraction of SVs without APH treatment are library artifacts vs. rare background events, but SVs accumulated above that background must have arisen during an experiment and provide a rich signal of dozens to hundreds of sequenced SV junctions per sample (**Figures 1C** and **1E**).

### SVs induced at CFSs by replication stress form preferentially in M-phase

A motivation in developing single-molecule svCapture was to determine when SVs form at CFS genes relative to replication. Replication stress occurs in S-phase, but we reasoned that subsequent junction formation could occur in S concurrent with fork failure, in G2, in M associated with MiDAS or other processes, or in the next G1 associated with 53BP1 foci ^41^. We therefore purified timed, flow-sorted cells and used real-time svCapture to assess when in the cell cycle APH-induced SV junctions formed.

Our first experimental paradigm matched that used to study MiDAS ^23^. HCT116 cells were treated with APH and synchronized at the G2-M boundary using the CDK1 inhibitor RO3306 (**Figure 2A**). Treatment timing helped ensure that arrested cells had experienced APH-induced replication stress in the prior S-phase. G2 (4N DNA content, phospho-histone H3 [pH3] negative) and M-phase (4N, pH3 positive) cells were harvested by flow cytometry prior to or 3h after release from RO3306 arrest, respectively (**Figures 2C** and **S3A** to **S3C**). svCapture revealed a small but statistically significant increase in deletion SV formation in G2 cells treated with APH in the prior S-phase (**Figure 2D**). However, deletion yield increased consistently by an average of 4.6-fold in M relative to G2-phase cells (**Figure 2D**). Because cells were held in colchicine and flow sorted, they could not have passed into the next G1, indicating preferential SV formation in M-phase. SVs that formed in M-phase had similar properties to those from bulk asynchronous cultures, including a large median size, a bias toward deletions, and a lower rate of induced duplications and inversions with distinct breakpoint distributions (**Figures S4A** to **S4C**).

**Figure 2.**
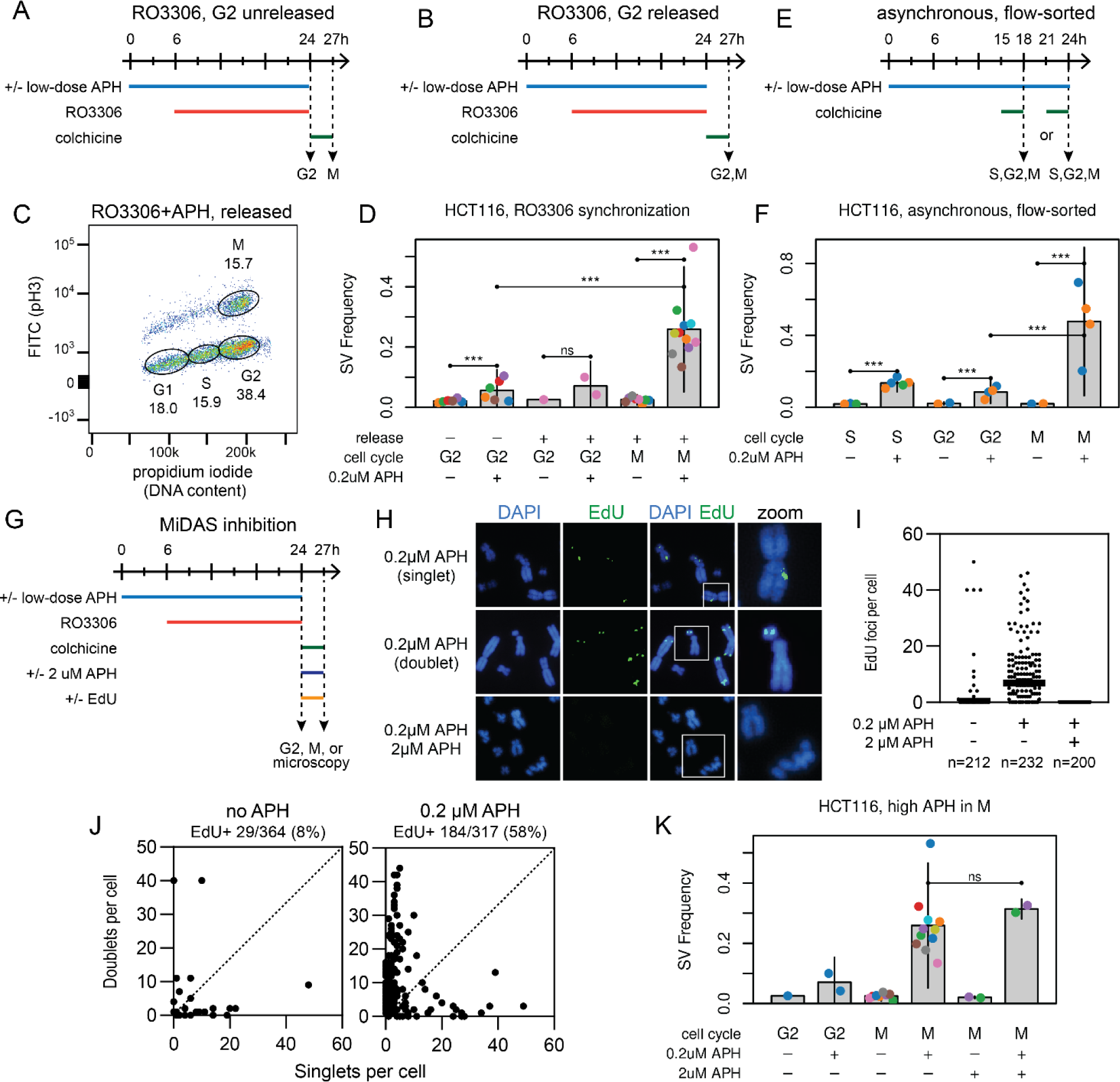
APH induces SV junction formation primarily during mitosis but independently of MiDAS. **A** and **B**. Two synchronization paradigms used to harvest APH-treated cells in different, timed phases of the cell cycle, where G2 cells were harvested before and after release from RO3306 arrest, respectively. **C**. Example flow sorting of HCT116 cells after release from RO3006. x-axis, DNA content; y-axis, pH3. **D.** SV frequencies from synchronized HCT116 cells harvested in the indicated cell cycle phases, showing increased APH-induced SV yield when cells passed from G2 to M. **E**. Timeline of experiments where cells in different cell cycle phases were flow sorted from asynchronous cultures. Colchicine improved M-phase cell yield and prevented re-entry into G1/S. **F.** SV frequencies from HCT116 cells using the paradigm in E, with M-phase cells again showing higher SV yield. **G**. Timeline of experiments where high-dose APH (2µM) was added at release from RO3306 to inhibit MiDAS. EdU was added at RO3306 release only when visualizing MiDAS foci. **H.** Example images of EdU foci visualized in M-phase showing singlet and doublet foci and the absence of foci with high-dose APH. **I.** Summary of EdU focus counts per cell for the indicated number of cells from two experiments. **J.** Comparison of EdU singlet and doublet focus yield between untreated and low-dose APH-treated cells over three experiments. Each point represents one cell with at least one EdU focus, stratified by its singlet (x-axis) vs. doublet (y-axis) focus count. **K.** SV frequencies from HCT116 cells harvested with and without high-dose APH suppression of MiDAS, which had no impact on SV yield.

To ensure RO3306 was not influencing results, *e.g.*, by altering replication dynamics or inhibiting replication-associated repair in G2 ^42, 43^, we modified the workflow by either (i) harvesting G2 cells after release from RO3306 (**Figure 2B**) or (ii) omitting RO3306 and using extended flow sorting to collect sufficient S, G2 and M-phase cells from asynchronous cultures (**Figure 2E**). In all cases, svCapture revealed a significant ∼5-fold increase in deletion SV yield in M relative to G2-phase (**Figures 2D** and **2F**). To explore whether G2 cells were less capable of SV formation because they were dying, we assessed cell death by monitoring cleaved caspase 3 relative to a positive control treated with etoposide (**Figure S3D**). G2-phase HCT116 cells did not show an excess of cleaved caspase 3 despite being substantially less likely to have formed SV junctions than pH3-positive M-phase cells.

### MiDAS is not required for SV formation at CFSs under replication stress

Preferential SV formation at CFSs in M-phase might suggest that SV junctions are created by error-prone MiDAS, which occurs preferentially in large, actively transcribed genes, including CFS genes ^23, 24, 25^. To test this possibility, we followed established protocols for inhibiting MiDAS by adding high-dose (2µM) APH upon release from RO3306 arrest (**Figure 2G**) ^23^. To ensure MiDAS suppression, we added EdU to parallel cultures after RO3306 release and examined M-phase-specific EdU foci by fluorescence microscopy (**Figure 2G**). High-dose APH abrogated MiDAS-associated EdU focus formation induced by low-dose APH (**Figures 2H-I**).

Interestingly, in contrast to some prior reports ^23^, most M-phase EdU foci induced by low-dose APH exposure in S-phase were doublets with signal on both chromatids consistent with semi-conservative DNA replication (**Figure 2J**), not the singlet foci restricted to one sister chromatid taken as evidence for conservative replication by MiDAS ^23^. Importantly, MiDAS inhibition by high-dose APH in M-phase HCT116 cells did not suppress deletion SV formation (**Figure 2K**), a finding supported by a single-replicate experiment in GM12878 cells (**Figures S4D**).

### MiDAS inhibition does not affect CFS expression in HCT116 or GM12878 cells

SV formation and CFS expression are different manifestations of replication stress at CFS loci ^18^. MiDAS inhibition was reported to greatly reduce total gaps and breaks and specific CFS expression in U2OS cells and to generate CFS-associated ultrafine bridges and increased nondisjunction of chromosomes 3 and 16, which contain the FRA3B and FRA6D loci, in U20S cells and MRC5 human fetal lung fibroblasts ^23^. Because we found no effect of MiDAS inhibition on FRA3B or FRA16D-associated SV formation in HCT116 or GM12878 cells, we explored the relationship between MiDAS, chromosome breakage, and specific CFS expression in those cell lines. We first determined the optimal timing of high-dose APH (2uM) for MiDAS inhibition as determined by EdU foci formation in M-phase in unsynchronized cultures (**Figure 3A**). In HCT116 cells, a 1-2h high-APH treatment before chromosome harvest eliminated MiDAS (**Figure 3B**). In contrast to HCT116 cells synchronized with RO3306 (**Figure 2H**), there were a small number of foci seen in cells treated with high APH for 3h, presumably representing cells entering mitosis form early G2 or S-phase (**Figure 3B**). For GM12878 cells, a 1h high-APH treatment eliminated MiDAS, with low levels of EdU foci appearing in cells treated for 2h (**Figure 3B**). Based on these results, we used use 1h and 2h high-APH treatments for cytogenetic analyses with both cell types. As with SV formation, we did not observe a significant difference in APH-induced CFS expression or in total gaps and breaks in either HCT116 or GM12878 cells upon MiDAS inhibition with high-dose APH (**Figure 3C** to **3F**).

**Figure 3.**
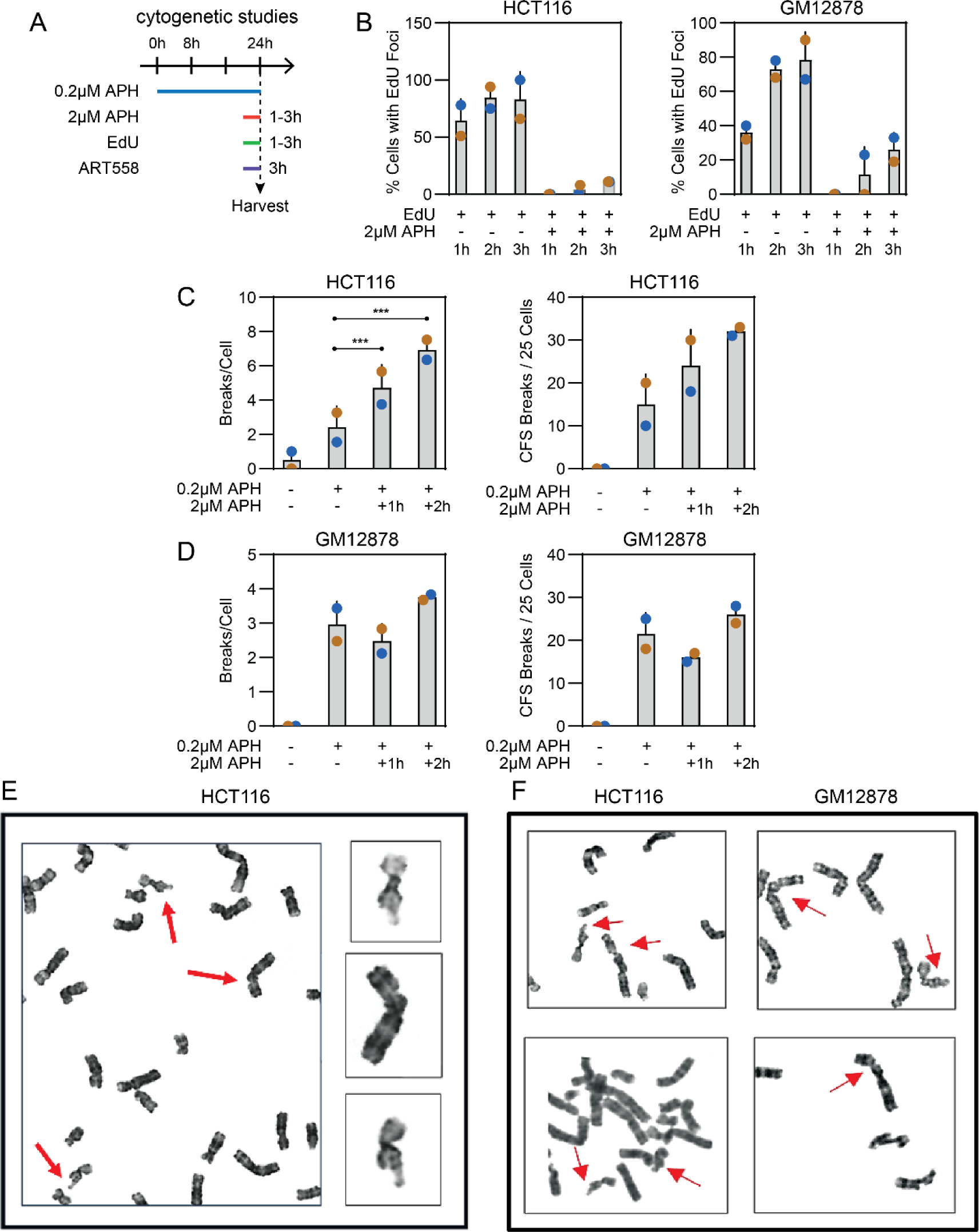
APH-induced chromosome gaps and breaks and CFS expression do not depend on MiDAS. **A**. Timeline of experiments where high-dose APH (2μM) and EdU were added at 1h, 2h, or 3h prior to chromosome harvest to inhibit MiDAS and for visualizing MiDAS foci, respectively. **B.** Comparison of EdU focus yield between untreated and high-dose APH-treated HCT116 and GM12878 cells. **C.** Total gaps and breaks and CFS expression in HCT116 cells with and without high-dose APH to suppress MiDAS; ***, p< 0.05. **D.** Like C, now for GM12878 cells. **E.** FRA3B and FRA116D CFS gaps/breaks (arrows) in a representative HCT116 cell treated with high-dose APH for 2h. **F.** Additional examples of FRA3B and FRA16D CFS gaps/breaks (arrows) in HCT116 and GM12878 cells treated with high-dose APH for 2h or 1h, respectively.

### SVs induced by replication stress have TMEJ-like junction profiles

Results above support preferential SV formation at large CFS genes in M-phase by a pathway other than MiDAS. To inform candidate alternative(s) we performed a detailed analysis of our extensive database of 12,786 *de novo* deletion junctions sequenced to base-pair resolution over all cells with unperturbed DNA repair. Short-read sequencing cannot reveal SVs created by HR, but prior microarray work never suggested events of that class ^17^. Instead, both microarray and svCapture data support junctions consistent with DSB end joining, where paired breakpoint positions in SV alleles revealed a range of microhomology usage, blunt joints, and *de novo* base insertions (**Figures 4A** to **4C**). The distribution of these junction classes, characterized by a prominent peak of 2bp microhomology and a substantial minority of *de novo* insertions, was strikingly reproducible across APH treatment conditions (**Figure 4C**), cell line and cell harvest workflow (**Figure S5A**), cell cycle phase (**Figure S5B**), SV type (**Figure S5C**), and MiDAS status (**Figure S5D**).

**Figure 4.**
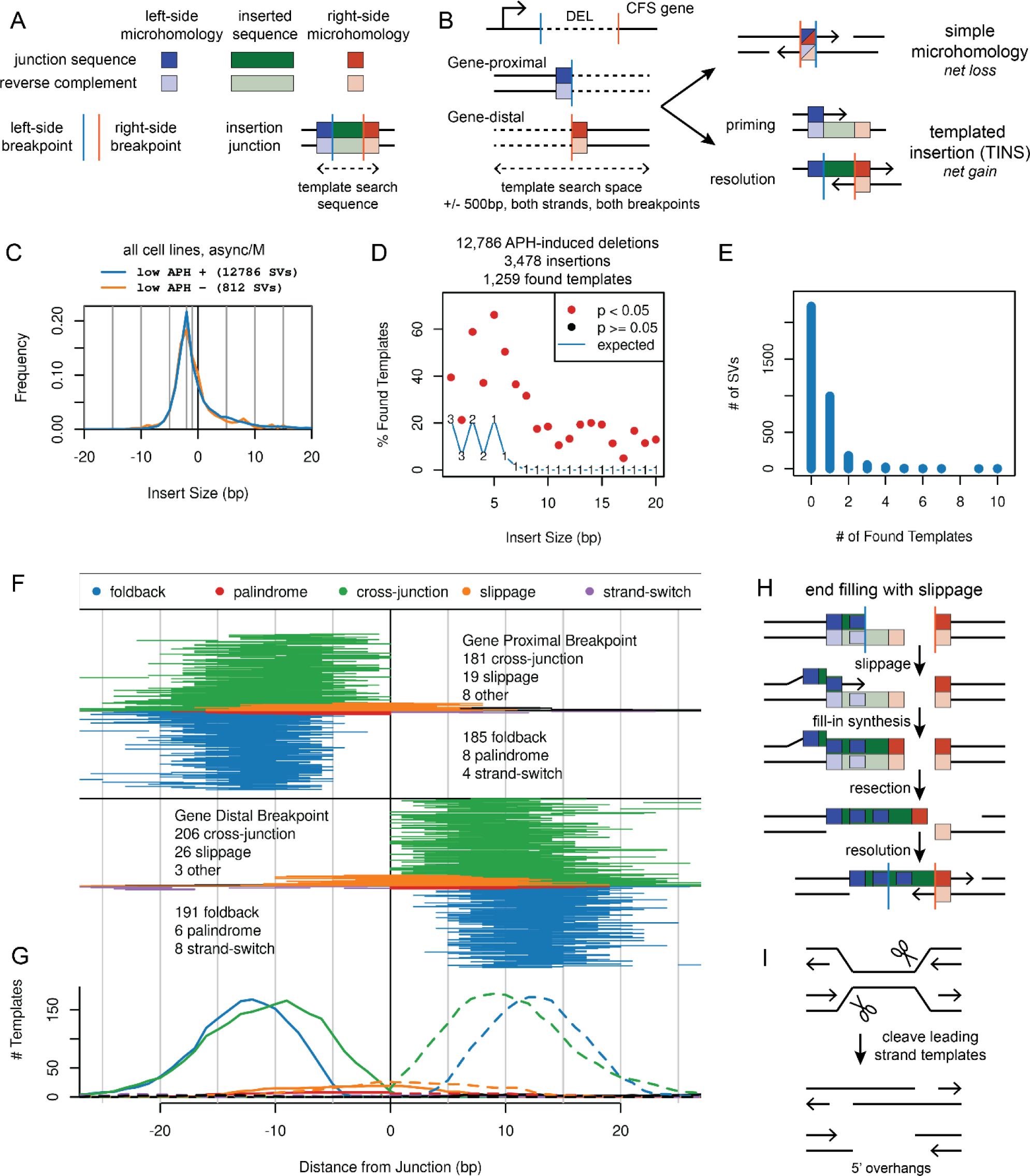
Junction analysis of >12K *de novo* deletions implicates TMEJ repair of DSBs created by leading strand cleavage. **A.** Guide to drawing conventions used in this figure. **B**. SVs are oriented to the transcription of the relevant CFS gene. Gene-proximal and gene-distal breakpoints can be paired via microhomologies that remove base pairs relative to the initial DSB ends, by blunt joints (not drawn), or by the insertion of novel bases, which might arise by template copying with sequential use of priming and resolving microhomologies. **C.** Distribution of net insert sizes from all cell lines harvested asynchronously or in M-phase, stratified by APH induction. Negative values represent microhomology usage. The peak is at - 2bp. **D**. Yield of identified templates for APH-induced deletions, stratified by insert size. To maintain specificity, smaller inserts required more bases of flanking microhomology (numbers on the random expectation line). **E.** Number of templates found for insertion SVs. Most had zero or one templates, supporting specificity. **F.** Pileup of insertion templates locations. The plot is oriented like panel B, *i.e.*, most templates were found in retained breakpoint segments. **G**. Histogram of the bases contributing to insertion templates, showing offset of foldback relative to cross-junction templates. Solid lines, left breakpoint; dashed lines, right breakpoint. Panels F and G share an X axis. **H.** Proposed end-filling slippage mechanism for some observed insertion events, where template bases cross into the deleted side of the inferred breakpoint position. **I.** Putative source of replication-associated, 5’ overhanging DSBs consistent with the slippage mechanism.

We characterized junction insertions in detail because they are a signature of TMEJ ^44, 45, 46^, a pathway recently implicated in mitotic DSB repair ^32, 33, 34^. Specifically, inserted bases are sometimes copied from template sequences near a DSB. The inferred repair process involves synthesis initiated from a priming microhomology flanking the inserted bases and eventual cross-DSB annealing via a second resolving microhomology on the opposite flank (**Figure 4B**) ^31, 45^. Three insertion classes with varying template orientations have been described, referred to here as foldback (also called inverse), cross-junction synthesis (also called direct), and a more complex and rarer strand-switching mechanism (**Figures S6A** to **S6C**) ^45^.

We searched for insertion templates 500bp upstream and downstream of each reference genome breakpoint (**Figure 4B**), requiring at least 7bp template spans to promoted specificity. We found a significantly higher fraction of templates than expected by random chance across all insertion sizes from 1 to 20 bp (**Figures 4D** and **4E**). However, we did not find templates for most insertions and the hit rate decreased with insertion size, possibly due to a correlated increase in the frequency of untemplated or multi-template events.

A highly informative footprint emerged from the 1,088 found templates featuring a peak of both priming and resolving microhomology lengths at 2 to 3bp (**Figure S5E**) and a net total template size, including flanking microhomologies, of typically less than 10 bp (**Figure S5F**). Templates were almost always found on the retained side of genomic SV breakpoints (**Figure 4F**), suggesting the search ensues after a DSB separates replicated/retained DNA from unreplicated/lost DNA. There was roughly equal utilization of foldback and cross-junction synthesis on either side of the SV junction, suggesting a random search (**Figure 4F**). However, the search was restricted in distance from the junction, with a strong peak of template bases within 20bp of breakpoints (**Figures 4F** and **4G**). Rare templates found at greater distances have a high likelihood of being random sequence matches (**Figure S7A**). The mechanistic specificity of templates near junctions was supported by a shift of foldback template bases nearest the junction toward positions further from the junction as compared to cross-junction synthesis (**Figures 4G** and **S7B**), consistent with the need for hairpin formation in foldback synthesis.

### A novel category of templated insertions suggests replication fork cleavage to 5’ DSBs

We observed two novel classes of insertion templates in our large number of *de novo* junctions. In the first, foldback synthesis appears to occur in a sequence that was likely coincidentally palindromic, given that the palindrome bases could not all anneal in the presumptive hairpin (**Figures 4F** and **S6D**). More notably, we observed templates on the top genome strand relative to a deletion that crossed from the retained into the lost portion of the corresponding genomic breakpoint (orange lines in **Figure 4F**). Importantly, designations of retained and lost DNA are relative to alignment breakpoint positions, not the source DSB termini whose structure we do not know. Thus, this insertion class can be modeled as end-filling events at putative 5’ overhanging DSB ends with slippage synthesis (**Figures 4H** and **S6E**). Strand melting and upstream reannealing of the 3’ terminus prior to end filling would lead to the discontinuity in genome alignment that causes DSB-terminal bases to be assigned as a *de novo* insertion past the apparent SV breakpoint. Slippage-class insertions have not been described for TMEJ occurring at CRISPR/Cas9-mediated DSBs ^31, 45^, likely because the mechanism invokes 5’ overhanging DSBs and thus does not apply to Cas9 blunt ends. In contrast, **Figure 4I** illustrates how cleavage of leading strand templates would lead to the DSB structure hypothesized to support slippage-class insertions.

### Chemical POLQ inhibition and POLQ knockout differentially impact SV formation

Our junction analysis implicated TMEJ as a possible mechanism of SV formation at large CFS genes. To test this hypothesis, we modified asynchronous and M-phase svCapture workflows to incorporate chemical POLQ inhibitors (**Figures 5A** and **5B**) and CRISPR-mediated *POLQ* knockout (KO) cell clones. We validated TMEJ loss using a published assay based on PCR detection of intracellular joining of transfected oligonucleotides (**Figures 5C** and **S8**) ^47^.

**Figure 5.**
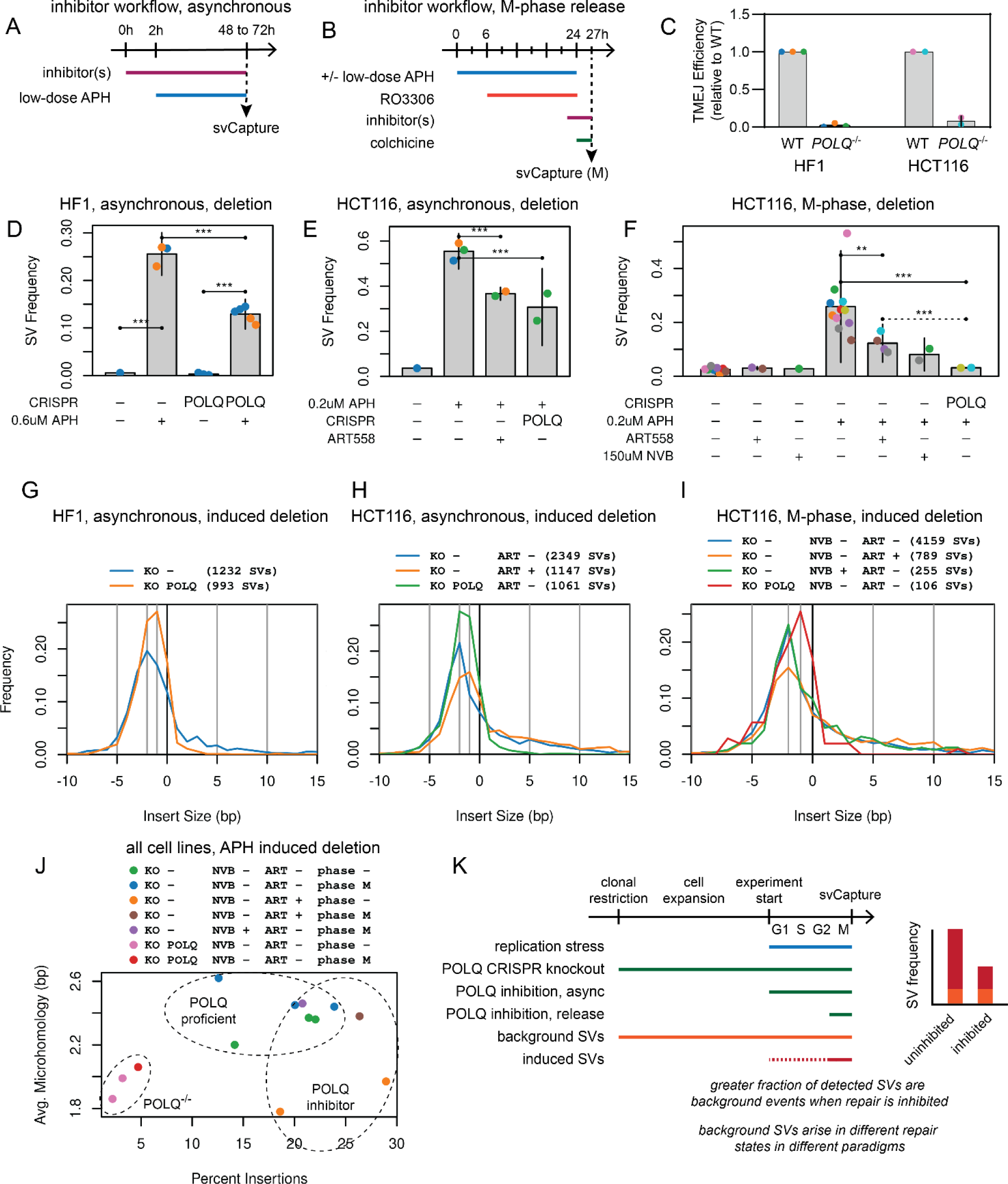
POLQ inhibition and knockdown reduce SV formation in mitosis. **A** and **B.** Modification of asynchronous and synchronization timelines, respectively, where DSB repair inhibitors were added prior to either APH addition or release into M-phase. **C.** Loss of TMEJ in *POLQ*^-/-^ HF1 and HCT116 cells as determined by joining of transfected oligonucleotides. **D** to **F.** SV frequency for the indicated cell lines, experimental paradigms, and mechanisms of POLQ suppression. All plots show deletion SVs. **G** to **I.** Insertion/microhomology size distributions for the data in D to F. **J.** Summary of junction property distributions. The x-axis is the percent of all SVs in a group that had 2 to 15 bp insertions, the y-axis is the average microhomology length of junctions without insertions. **K.** Differences between repair suppression paradigms with respect to the SV junctions that were ultimately sequenced.

Results consistently supported a role for POLQ in SV formation at CFSs but varied by cell type and method with different degrees of SV loss. Asynchronous *POLQ*^-/-^ HF1 and HCT116 cells each showed a reproducible, significant, but partial loss of deletions as well as other types of SVs relative to wild-type (**Figures 5D, 5E**, **S9A** and **S9B**). Deletion SV reduction was again partial in HCT116 asynchronous or M-phase cells treated with the POLQ inhibitors ART558 ^48^ or novobiocin (NVB) ^49, 50^ (**Figures 5E** and **5F**). In contrast, APH-treated *POLQ*^-/-^ HCT116 M-phase cells showed baseline levels of SV formation with no apparent induction by APH (**Figures 5F** and **S9C**).

Because SV frequency alone might not reveal the full role of TMEJ if another repair pathway could partially replace it, we analyzed properties of residual SVs from cells with impaired TMEJ. SV sizes were similar regardless of POLQ status (**Figures S9D** to **S9F**). In contrast, we observed shifts in junction distributions comprising changes in microhomology lengths and insertion frequencies. *POLQ*^-/-^ cells across all cell lines and workflows showed a near absence of insertions >=3bp and a shift in peak microhomology length from 2bp to 1bp (**Figures 5G** to **5I** and **S9G** to **S9I**). In contrast, POLQ chemical inhibition did not substantially reduce insertions while the shift toward shorter microhomologies sometimes remained apparent (**Figures 5H** to **5I** and **S9H** to **S9I**). We found templates for insertions detected in ART558 and NVB-treated samples much like uninhibited cells, although with a higher relative rate of slippage insertions (**Figure S10**). These results were supported by a single experimental replicate in GM12878 cells (**Figures S9J** to **S9L**).

Figure 5J summarizes how POLQ-proficient, POLQ-inhibited, and *POLQ*^-/-^ samples group with respect to insertion frequency and microhomology lengths. The dynamics of these different manipulations must be carefully considered when interpreting results (Figure 5K). POLQ protein loss is distinct from its chemical inhibition, which may only partially inhibit enzymatic activity and/or permit structural roles to be fulfilled. Moreover, inhibitors were used transiently whereas KO cells lacked POLQ from the time they were cloned. SVs that arose as background events before an experiment would form in POLQ proficient vs. deficient states for inhibition vs. KO, respectively, and background events become a larger fraction of detected SVs as APH induction decreases (Figure 5K).

### TMEJ and NHEJ cooperate in SV formation in some asynchronous cells

Despite challenges comparing POLQ chemical inhibitors and KO clones, results above demonstrate that some SV formation at CFSs can occur without POLQ, especially in asynchronous cultures. To explore the interplay between TMEJ and NHEJ in different cell cycle stages, we added chemical inhibition of DNA-PKcs using NU7441 ^51^ and CRISPR/Cas9 KO of DNA ligase IV gene *LIG4* (**Figure S11A** and **S11B**) ^52, 53^. svCapture deletion yield in asynchronous HF1 cells was not altered by NU7441 in either wild-type or *POLQ*^-/-^ backgrounds (Figure 6A). In contrast, NU7441 significantly decreased deletion yield relative to ART558 or *POLQ* KO in asynchronous HCT116 cells (Figure 6B) and in GM12878 cells in a single replicate (**Figure S11C**). This synergy was especially apparent when ART558 was added to asynchronous cultures of *LIG4*^-/-^ HCT116 cells, which abrogated APH-induced SV formation (Figure 6B).

**Figure 6.**
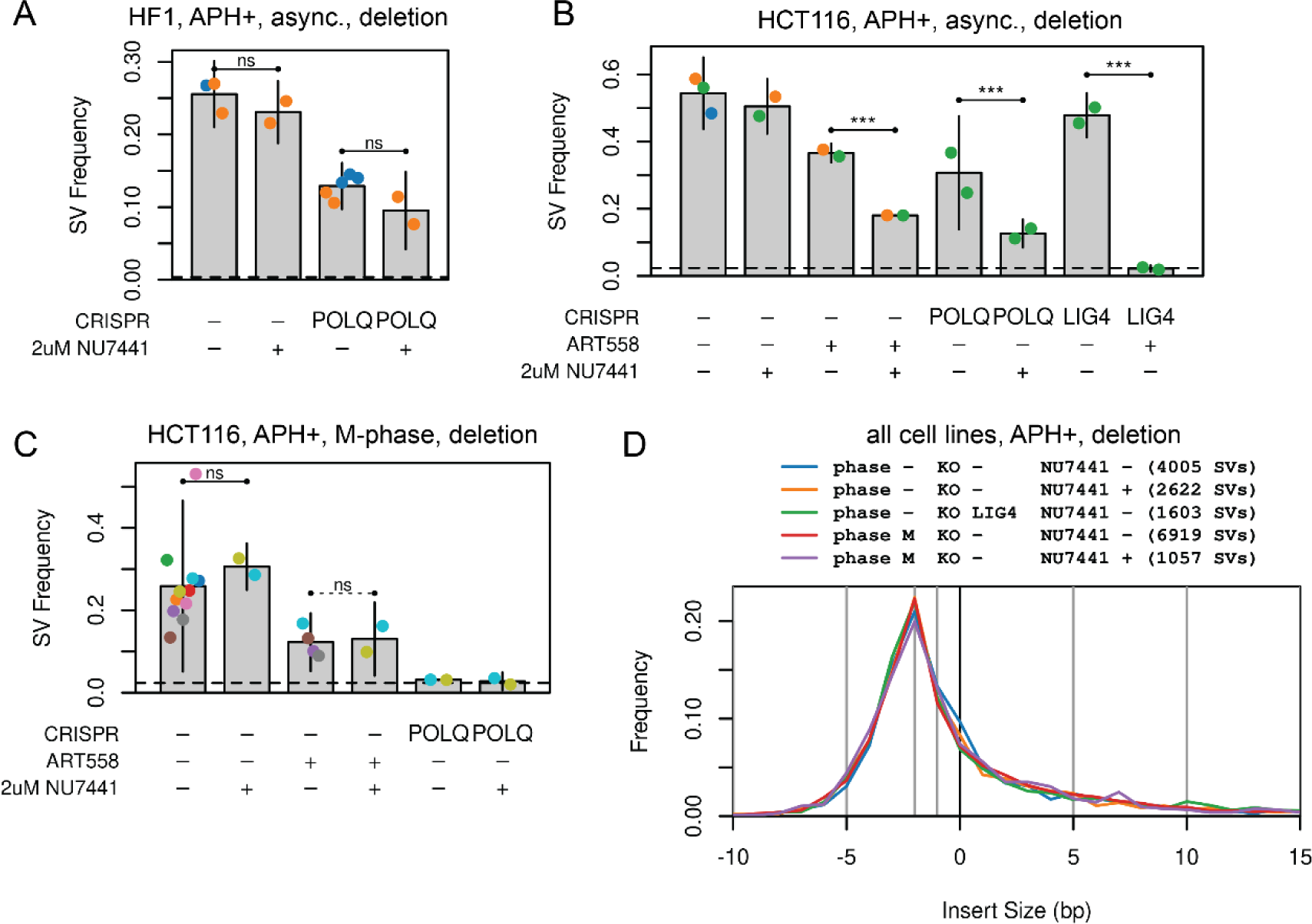
Cell-cycle dependent interplay between TMEJ and NHEJ in SV formation at CFSs. **B.** Deletion SV frequency in asynchronous, APH-induced HF1 cells as a function of *POLQ* knockout and NHEJ inhibition with Nu7441. **C.** Deletion SV frequency in asynchronous, APH-induced HCT116 cells as a function of *POLQ* and *LIG4* knockout, and inhibition of NHEJ with Nu7441 or TMEJ with ART558. **D.** Like A and B, now for HCT116 cells released into M-phase following RO3306 arrest. For clarity, panels A to C only show samples induced to form SVs with low-dose APH. Horizontal dashed lines indicate the cell-line-specific SV level without APH addition. **E.** Like Figure 4C, showing no effect of NHEJ loss on junction microhomology and insertion profiles.

A possible confounder above is that loss of both TMEJ and NHEJ can impair cell growth in some contexts ^54^, although we did not observe excessive cell death. To restrict the timeframe during which cells were doubly deficient, we added NU7441 to M-phase HCT116 cells just before release from RO3306 and observed that NU7441 now had no incremental impact on SV formation (Figure 6C). As noted above, *POLQ* KO alone was sufficient to abrogate APH-induced SV formation in M-phase HCT116 cells (Figure 6C). Throughout, loss of NHEJ by either chemical inhibition or *LIG4* KO had no impact on junction insertion/microhomology distributions or insertion template locations (Figure 6D and **S11D**), consistent with these baseline junction properties being driven by TMEJ. Interestingly, NU7441 also had no impact on junction insertion/microhomology distributions in *POLQ*^-/-^ cells (**Figure S11E**).

## Discussion

Genomic SVs and CNVs are a major cause of genomic imbalance in the germline, normal tissues, and in cancer cells. This study addresses nonrecurrent SVs that arise in non-repetitive loci by mechanisms linked to negative interactions between transcription and replication. Unlike the early replicating genome, where these interactions include machinery collisions and R-loop formation ^22, 55, 56, 57^, genomic instability in the late replicating genome relates more to replication-transcription conflicts that cause unreplicated DNA to be propagated into M-phase ^23, 24, 25^. The most unstable late-replicating loci are at the largest human CFS genes, where transcription is required for instability by impeding origin usage and completion of replication ^22, 40, 58^. We show that some of the mechanisms evolved to resolve the resulting unreplicated DNA prior to cell division can create large non-recurrent SVs.

We used new genomic technologies to monitor SV junctions as they formed in cultured cells. svCapture ^35^ was highly effective at detecting *de novo* SV junction formation in part due to the very high rate of CNV formation that occurs at transcribed CFS genes under replication stress ^17^. We characterized 45,507 independent *de novo* SVs in controlled, prospective experiments with a combined 282,763-fold target region coverage, which provides a uniquely powerful dataset for exploring junction formation mechanisms. Multiple observations affirm that we are measuring *bona fide* SV junctions at CFSs. All aspects of CFS CNVs observed in prior microarray studies ^12, 13, 14, 15, 16, 17, 18^ were recapitulated by svCapture, but with much greater resolution and data density. These included APH induction, a strong bias toward deletions despite a clear ability to detect duplications and inversions, the wide distribution of junctions within targeted loci that matches CNV microarray breakpoints, and the greater tendency of duplications to flank the central CFS region. Moreover, distributions of junction microhomology and templated insertions were pathognomonic of cellular DNA repair capacity and must therefore primarily comprise SVs created in cells.

Mathematical models ^59^ have shown that stochastic replication fork failure coupled with an inability to rescue replication via dormant origins can alone yield the diffuse central peak of SV junctions we see within CFS genes ^17^. Consistently, we found no subregions in any of the CFS genes we studied that acted as more highly localized SV hotspots, despite having targeted 250kb or 400kb of different genes. The large introns in CFS genes often have AT-rich sequences that have been invoked as important for CFS expression through the formation of focal flexibility peaks ^37, 38^. Biochemical studies of DNA polymerase progression through specific focal regions of CFS genes, especially sequences in *WWOX*, further showed that they can impose mechanistic barriers to replication ^60, 61^. However, no focal regions could be correlated with SV junction locations in our svCapture data. The mechanisms leading to SV formation at CFS genes link transcription to a diffuse distribution of *de novo* junctions. A limitation making conclusions about the lesions giving rise to those SVs is that extensive processing of DSBs, whose location and structure we cannot determine from junction sequences alone, would make the SV distribution appear more diffuse than that of the source lesions.

CFSs replicate as late as M-phase ^17, 20, 21, 23, 62^, and the distribution of SV junctions we have observed at CFS genes corresponds well to the location of MiDAS synthesis peaks observed under replication stress ^24, 25^. Thus, replication inhibition and the initiation of instability begin in S-phase but are not resolved until completion of replication late in the cell cycle. We hypothesized SV formation would also occur in late G2 or M-phase coincident with completion of replication. This hypothesis was confirmed by cell-cycle timing experiments that showed by direct measurement that 80% of CFS SV junction formation occurs after cells pass into M-phase. Specifically, most SV junction formation occurred after the appearance of pH3 in chromatin coincident with MiDAS timing as previously reported ^23^. Experimental variations indicated that RO3306 was not artifactually suppressing SV formation in G2-phase secondary to CDK1 inhibition ^42, 43^. The use of colchicine and flow sorting further ensured that junctions were not formed secondary to rupture of ultrafine bridges and progression into the next G1 where 53BP1 bodies result ^23, 41^, although the latter pathway remains an interesting additional unexplored possibility.

Describing M-phase SV formation depended on the single-molecule sensitivity of svCapture, since nonrecurrent SV junctions will only be present in a single copy until replicated in the next cell cycle. Prior work examined features such as single-chromosome end-to-end fusions in Ku-deficient cells, radial formation in Fanconi anemia cells, and the persistence of DNA damage evident as DNA repair foci ^33, 63^, but to our knowledge ours is the first direct demonstration of preferential intrachromosomal SV junction formation in M-phase.

Within M-phase, multiple possibilities existed for SV junction formation. Because MiDAS is an error-prone form of conservative replication ^23, 26, 64^, and because MiDAS hotspots are found in CFS loci ^23, 24, 25^, it was a plausible mechanism for executing SV junction formation. However, MiDAS proved to not be required nor did suppressing it significantly increase SV junction formation. MiDAS suppression also did not block or reduce CFS expression in GM12878 or HCT116 cells, contrary to prior studies with other cell types ^23^. Thus, MiDAS appears to be a genome preserving pathway for completing synthesis of unreplicated DNA in mitosis, but for reasons that are not fully clear, MiDAS and SV formation do not appear to function as competing pathways, *i.e.*, with one increasing as the other decreases. Importantly, the spans of unreplicated DNA passing into M-phase at CFS genes are exceptionally large relative to expectations for the rest of the genome under more typical cellular stresses. MiDAS may never be fully effective in completing replication of hundreds of kb of CFS DNA, which might explain our data if SV formation could occur even after partial MiDAS.

Our first evidence that TMEJ mediates M-phase SV formation came from detailed analysis of 31,497 fully sequenced SV junctions, which showed a remarkably reproducible pattern across SV types, cell cycle phases, and APH-induction status. In repair-proficient cells, that profile included a prominent peak at 2bp microhomology and a long tail of ∼20% *de novo* base insertions, a signature of TMEJ ^45, 46, 65^. Short-read sequencing has detection biases against longer insertions >20bp but is fully able to reveal nonhomologous junctions with several bases of microhomology or insertion. Thus, the junction profile we observed is a reliable signature for the mechanism(s) catalyzing *de novo* SV junction formation, especially in M-phase. SV junctions arising at CFS genes used insertion templates restricted to a window of approximately 20bp in the retained/replicated segments surrounding the breakpoint positions in the source loci. These insertion events were lost in *POLQ*^-/-^ cells supporting their formation by TMEJ and establishing constraining parameters against which biochemical and structural studies of POLQ should be compared. Interestingly, templates were found for a minority of *de novo* insertions. We suspect this is mostly due to iterative copying of multiple templates and/or untemplated synthesis by POLQ ^65, 66^ but cannot rule out other processes. Unfortunately, the small size of insertions prevents genome-wide searches for possible templates.

The most definitive demonstration of a primary role for POLQ in SV junction formation was the loss of APH-induced M-phase SVs in *POLQ*^-/-^ HCT116 cells. That result is strongly consistent with recent studies identifying POLQ suppression by RAD52 and BRCA2 prior to M-phase ^67^ and its activation in M-phase by RHINO-mediated recruitment to DSBs and phosphorylation by PLK1 ^32, 33, 34^. Our results support the idea that TMEJ acts an error-prone rescue pathway for dealing with unreplicated DNA in M-phase to prevent mitotic catastrophe. TMEJ has been shown to produce short 50-200bp deletions in mutation accumulation experiments in *C. elegans* ^68^ and 5-50bp deletions at Cas9-induced DSBs in mouse cells ^69^ but those small SVs are best modeled as occurring via resection of a single DSB. To our knowledge, ours is the first experimental demonstration that POLQ mediates the formation of large, multi-lesion, spontaneous SVs >100kb in mammalian cells, and that it occurs mainly in M-phase.

Given the above, we were surprised by the partial effects of POLQ inhibitors on SV formation, mainly the widely used and more specific agent ART558. Doubling the concentration used from 5µM to 10 µM, concentrations used by others ^48, 70^, gave similar results in multiple cell lines. POLQ inhibition did sometimes reduce microhomology lengths at SV junctions similarly to *POLQ* KO, arguing we effectively inhibited POLQ, but with a relative persistence of templated insertions. Because our experiments were based on SV junction formation as the primary measured outcome, they may reveal and support a separation of POLQ functions ^71^.

ART558 binds to the polymerase catalytic domain of PolQ and inhibits its activity ^48^, whereas NVB inhibits the POLQ ATPase ^49, 50^. Neither removes the potential for POLQ acting in non-catalytic ways. Based on our results, microhomology use appears to be more impacted by alterations in POLQ catalytic activities whereas insertion formation strongly depends on structural functions. However, chemical inhibition of POLQ was transient whereas POLQ KO preceded the experimental window, which impacts the SVs that svCapture would detect as background events.

Despite clear evidence that TMEJ acts in M-phase as a major source of large SVs, that association is incomplete. Even with *POLQ* KO, some *de novo* SV formation remained in asynchronous cell cultures. This effect could not be explained by different M-phase behavior of cancerous and normal cells ^72^ because it was observed in both the HF1 and HCT116 cell lines. These results suggest that some SV formation can be catalyzed by another end joining pathway, at least as a backup to TMEJ. Importantly, in both the current experiments and in our prior study of murine Xrcc4^-/-^embryonic stem cells ^73^, we saw no impact of the loss of NHEJ alone on SV formation, identifying it as secondary to TMEJ. It is difficult to rule out that some SVs detected in S or G2 were formed in a prior M-phase but given prior knowledge that NHEJ is most active in G1 ^11^ and thought to be largely inactive in M-phase ^74, 75^, it seems more likely that non-TMEJ pathways had a greater role in SV formation outside of M-phase. Of note, DNA polymerase lambda has been suggested to mediate end joining independently of NHEJ and TMEJ and might play a role in SV formation ^76^.

Insertion templates that appeared to cross breakpoint junctions were unexpected but can be modeled as arising from polymerase slippage during end-filling of 5’ overhanging DSBs, presumably DSBs created in M-phase by fork cleavage that activates replication rescue. Both the MUS81 and GEN1 structure-specific nucleases are required for CFS expression ^77, 78^ and are excellent candidates for creating those source DSBs leading to SV formation. If our slippage model is correct, it implicates MUS81 as the primary nuclease as it is thought to cleave the leading strand template at stalled forks, the orientation that would yield 5’ overhangs ^79^. Such symmetrical cleavage of stalled forks has been proposed to lead to HR-independent SCEs ^80^ where DSB repair by TMEJ would obligatorily lead to SV formation, especially deletions corresponding to unreplicated DNA spans. It is less obvious how TMEJ would lead to duplication and inversion SVs, which, although less frequent, were seen at CFSs and bore the same hallmarks of TMEJ.

Our results provide key insights into the ways that error-prone replication rescue in M-phase proceeds from DSB formation to the creation of various types of SVs by TMEJ. It is not yet known to what extent the final resolution of replication-associated damage throughout the genome is deferred until M-phase, but MiDAS-associated propagation of unreplicated, R-loop-associated DNA into M-phase has been observed in *BRCA2* and *RAD51*-deficient cells and cells with cyclin E1 overexpression ^56, 81, 82^. Moreover, we have observed replication stress-induced CNVs that mimic mammalian nonrecurrent CNVs at numerous non-CFS loci using microarrays^17^. Taken together, it seems likely that SVs in non-CFS loci may also be created in M-phase by TMEJ. Indeed, our results closely match prior descriptions of microhomologies and frequent insertions at SV junctions in both normal human CNVs ^46, 83, 84^ and SVs in cancers ^85, 86, 87^, strongly implicating POLQ and mitotic TMEJ as central mechanisms in mammalian SV mutagenesis. Further work is needed to understand the endogenous or physiologically relevant exogenous stressors leading to M-phase SV formation and to reveal the precise nature of the lesions that are repaired to SV junctions in M-phase.

## Methods

### Cell Culture Models

#### UM-HF1 fibroblasts

UM-HF1 (abbreviated HF1 throughout) is a XY male, euploid, TERT-immortalized human foreskin-derived fibroblast cell line derived and maintained at the University of Michigan ^17^ subject to human data access restrictions. It has known CFSs/CNV hotspots at genes *PRKG1*, *NEGR1*, and *MAGI2* that provide excellent svCapture signal in a non-cancerous cell line ^17, 35^, but HF1 cells are not easily synchronized for cell-cycle analysis. HF1 cells were cultured at 37C with 5% CO2 in Dulbecco’s Modified Eagle Medium supplemented with 13% fetal bovine serum (FBS), 2mM L-glutamine, and 100 U/ml penicillin-streptomycin (Gibco).

#### GM12878 lymphoblastoid cells

GM12878, (Coriell, RRID CVCL_7526), is a highly studied XX female, euploid, EBV-immortalized human lymphoblastoid cell line generated as part of the HapMap Project. It has CFSs common to lymphoblastoid cells at genes *WWOX* and *FHIT* that allow direct comparison of CFSs and SVs in a suspension cell line. GM12878 cells were cultured at 37C with 5% CO2 in RPMI 1640 medium supplemented with 15% FBS, 2mM L-glutamine, and 100 U/ml penicillin-streptomycin.

#### HCT116 colon cancer cells

HCT116 (ATCC, RRID CVCL_0291) is a highly studied male, mismatch-repair deficient, human colon cancer cell line. This work established that genes *WWOX* and *FHIT* are SV hotspots in HCT116, consistent with CFS expression analysis, gene expression analysis using Bru-seq ^88^, and genomic analysis that showed baseline SVs in these genes ^89^. HCT116 cells were cultured at 37C with 5% CO2 in McCoy’s 5A medium supplemented with 10% FBS, 2mM L-glutamine, and 100 U/ml penicillin-streptomycin.

#### CRISPR-Cas9-mediated gene knockout

For CRISPR-Cas9 mediated knockout (KO) of *POLQ* or *LIG4* in HCT116 cells, single guide RNAs (sgRNAs, **Table S4**) were designed using CHOPCHOP ^90^ and cloned into the PX459 plasmid, which carries the human U6 promoter, Cas9 gene, and puromycin resistance gene ^91^. The plasmids were transfected into HCT116 cells using Lipofectamine 3000 (Invitrogen). For *POLQ* KO in HF1 fibroblasts, vector pLentiCRISPR v2 with integrated sgRNAs (GenScript) was transfected into HF1 cells by the University of Michigan Vector Core. Following transfection, cells were subjected to selection with 1µg/ml puromyocin and KO clones were established by plating at low-density and isolating single colonies with cloning rings for expansion in multi-well dishes. Following clonal expansion, PCR with primers flanking the sgRNA binding sites was used to amplify the target alleles followed by Sanger sequencing. The resulting mixed allelic sequence traces were analyzed using Synthego ICE software ^92^. Clones were preferred for further use when each of the two alleles yielded distinct frameshift mutations (**Table S4**). Clones that yielded identical biallelic mutations were confirmed with ddPCR to establish a copy number of two for the mutant allele. *LIG4* KO clones were further validated with immunoblotting. The large protein size and low expression of POLQ resulted in inconclusive westerns, so *POLQ* KO clones were validated using the TMEJ assay described below. At least two independent clones of all mutated cell lines were frozen at early passage post-cloning for subsequent SV analysis.

#### Replication stress induction and monitoring

The DNA polymerase inhibitor aphidicolin (APH, Sigma) was dissolved in DMSO at a stock concentration of 200µM. For CNV and CFS induction, cells were cultured with APH as follows: HF1, 0.6 µM; GM12878, 0.4µM; HCT116, 0.2µM. These doses were empirically determined per cell line to be consistent with slowed but continued cell division and to produce approximately 2-5 chromosome gaps and breaks per cell in GM12878 cells. These concentrations of APH are defined as low-dose APH throughout. The duration of low-dose APH treatment and its timing relative to other cell manipulations varies is shown in timelines in relevant figures. High-dose APH (2µM) and EdU (10µM), to suppress MiDAS or reveal MiDAS foci, respectively, were added 1 to 3h prior to harvest.

#### Chromosome breaks and common fragile sites

CFSs and total gaps and breaks were scored on Giemsa-banded metaphase preparations following 24h low-dose APH induction with or without the addition of high-dose APH as described above. Cells were harvested for chromosome preparations using standard conditions of a 20 to 45min Colcemid treatment (50ng/ml: Gibco) followed by a 15min incubation in 0.075M KCl hypotonic solution at 37C and multiple changes of Carnoy’s fixative (3:1 methanol:acetic acid). Fixed cells were dropped onto glass slides to generate metaphase spreads and slides were baked overnight at 60C before Giemsa banding. For Giemsa banding, slides were dipped in water, treated with trypsin solution (0.0005% trypsin and 0.02% Tyrode’s diluted in HBSS) for 50s, followed by two rinses with 0.9% NaCl, stained in Giemsa staining solution (5% Giemsa in Gurr Buffer, pH 6.8) for 5min, followed by two sequential rinses in water. Metaphase chromosomes were visualized using Zeiss Axiphot microscope and chromosome breaks and gaps were analyzed in 25-50 metaphases from each experimental sample.

#### Cell synchronization and flow sorting

Cells were treated with low-dose APH for an initial 6h and then arrested at the G2/M boundary by addition of 9µM (HCT116) or 10µM (GM12878) RO3306 (ApexBio) with continued APH for an additional 18h. Parallel cultures were then either harvested for flow sorting of S and G2 fractions or washed three times with PBS and released into warm media containing 75ng/ml colchicine for 3h for flow sorting of G2 and M fractions, and also with 10µM EdU for cells used to visualize MiDAS foci. For cells treated with novobiocin (NVB, 150µM, Sigma), the drug was added together with low-dose APH and added back again after release from RO3306. ART558 (5µM or 10µM, MedChemExpress) or Nu7741 (2µM, Fisher Scientific) were added 2h prior to harvest for S and G2 fractions or RO3306 release and again added back to the media after release. High-dose (2µM) APH was added for 3h after cells were released from RO3306. When performing cell cycle analysis without RO3306, asynchronous HCT116 cells were treated for 24h with low-dose aphidicolin. Three hours prior to harvest 100ng/ml Colcemid was added to the media to enrich the M-phase population.

For flow cytometry, cells were harvested with trypsinization, collected in cold media, spun down (5min, 500xg, 4C), and fixed in 70% ethanol overnight at −20C at a concentration of 1×10^6^ to 2×10^6^ cells/ml. Cells were then washed with PBS, permeabilized with 0.25% Triton X-100 in PBS on ice for 15min, spun down, and stained with phospho-histone H3 (pH3) antibody (Cell Signaling Technology) conjugated to Alexa fluor 488 (Invitrogen) at a 1:50 dilution in antibody staining buffer (0.5% bovine serum albumin (BSA) in PBS) for 1h. Cells were washed twice with antibody staining buffer and stained with a solution of 100µg/ml propidium iodide and 100µg/ml RNAse. Samples were then submitted to the University of Michigan Flow Cytometry Core for collecting cell cycle fractions using a FACS Aria III (BD Bioscience) or Bigfoot Cell Sorter (ThermoFisher). Gating established that S-phase fractions had a DNA content between 2N and 4N and were pH3 negative, G2-phase fractions had 4N DNA content and were pH3 negative, and M-phase fractions had 4N DNA content and were pH3 positive. Flow sorting was continued until at least 200,000 cells had been collected from all target cell cycle fractions in a sample.

#### MiDAS assessment

After initial cell harvesting, cells treated with EdU after release from RO3306 were prepared for metaphase analysis as described above for chromosome spreads. MiDAS activity was then assessed using Click-iT reaction and Alexa Fluor 488 azide (Invitrogen). Slides were first treated with 4% formaldehyde in PBS for 4min, washed three times with PBS, and blocked with 3% BSA in PBS for 30min. Permeabilization and the Click-iT reaction were performed according to the manufacturer’s instructions. Slides were then washed with 3% BSA/0.5% Triton X-100 in PBS three times for 10min per wash, rinsed with water and mounted with Prolong Gold DAPI Antifade mounting media (Sigma). Metaphase chromosomes were visualized using Zeiss Axiphot fluorescence microscope. Images were acquired using CellSens software. EdU quantification was done manually at the microscope, counting for every cell the number of EdU foci stratified by whether just one (singlet) or both (doublet) chromatids were labeled.

#### TMEJ and NHEJ assays

To monitor the efficacy of chemical and genetic interventions intended to inhibit POLQ/TMEJ or NHEJ, we used an assay based on transfected oligonucleotides substrates subjected to intracellular end joining ^47^. The substrates were prepared by the Dale Ramsden laboratory by annealing substrates in a buffer containing 10mM Tris-HCL, pH 7.5, 100mM NaCl, and 0.1 mM EDTA. 5ng of NHEJ substrate or 500ng of TMEJ substrate were then electroporated separately in a solution containing 500ng pUC19 plasmid, 0.16µl 2X PBS, 0.84µl EB buffer and buffer R in a total of 10µl into 200,000 cells using the Neon system with a single pulse of 1,350V (GM12878) or 1,530V (HCT116) for 20ms. Prior to electroporation, cells were pretreated for 2h with 10µM ART558, 150µM NVB, or 2uM Nu7441. After electroporation, cells were incubated in antibiotic-free media supplemented with the appropriate drug for another 30min at 37C, followed by a wash with PBS, and incubated at 37C for 15min in 40µl HBSS containing 125U Benzonase and 5mM magnesium chloride. DNA was extracted using the QIAamp DNA mini kit (Qiagen) following the manufacturer’s protocol with the addition of 1mM EDTA added to buffer ATL. Samples were then analyzed using qPCR with TaqMan Fast Advanced Master Mix primers and probes using 7500 Real-Time PCR System (Applied Biosystems). The cycling conditions were 50C for 2min, then 95C for 2 min, followed by 40 cycles of 15s at 95C and 1min at 60C.

For *POLQ* CRISPR KO, samples were normalized to the NHEJ substrate whereas *LIG4* CRISPR KO samples were normalized to the TMEJ substrate. Equal amounts of DNA were used for samples treated with inhibitors. The wild-type or untreated samples were used as reference to calculate ΔΔCt values.

#### Apoptosis via cleaved caspase

To assess cell death that might result from cell treatments above, we monitored cleaved caspase-3, a marker for apoptosis. Cells were treated and prepared as described above for synchronization and flow cytometry with the addition of cleaved caspase-3 antibody (Cell Signaling Technology) conjugated to Alexa Fluor 647 added together with pH3 antibody. As a positive control for apoptosis, cells were separately treated with 10µM etoposide (Sigma) for 72 hrs. Data were assessed for the percentage of cells positive for cleaved caspase-3.

#### svCapture library preparation and sequencing

At least 200,000 cells were collected and centrifuged for 10 min at 1000 rpm from either bulk asynchronous cultures or flow-sorted cell-cycle phases. Supernatant was removed until 100µl remained and 200µl DNA/RNA shield (Zymo Research) was added with 15µl 20 mg/ml proteinase K and incubated for 20min at room temperature. Genomic DNA was purified using Quick-DNA microprep plus kit according to the manufacturer’s instructions (Qiagen). Further processing steps through high-throughput sequencing were performed at the University of Michigan Advanced Genomics Core. Bead-based tagmentation libraries were prepared with the Illumina DNA Prep with enrichment kit, using 300ng of genomic DNA, IDT for Illumina unique dual barcodes, and library PCR amplification of nine cycles. Libraries were quantified using Qubit and quality checked using an Agilent TapeStation to ensure that at least 350ng and preferably 500ng of prepared library was available to support robust target capture.

Hybridization capture probes were targeted to the central 250kb or 400kb of large CFS genes, the region of peak accumulation of SV breakpoints ^17^. Final probes were designed to be target-specific and synthesized by Twist Biosciences using their proprietary algorithms and used as provided by the vendor. Capture was performed by pooling 500ng of each library and hybridizing with 4μl Twist Biosciences probes and 6μl PCR grade water. Target enrichment on magnetic beads was performed according to manufacturer instructions. Retained library fragments were amplified with 12 cycles of PCR for sequencing.

Sequencing reads were obtained in the 2 × 151 format using Illumina NovaSeq 6000 or Illumina NovaSeq X Plus. Barcoded samples were pooled and subjected to a sequencing depth calculated to yield a projected coverage of ∼2,000-fold in the capture target regions based on experience ^35^, typically 2.5% of a NovaSeq S4 flow cell per sample. Insert size and target region coverage were maintained over narrow ranges over all analyzed samples (**Figure S1B**).

### svCapture data analysis

#### svCapture pipeline execution

We previously reported the svCapture data analysis pipeline and Shiny app ^35^ constructed in the svx-mdi-tools suite of the Michigan Data Interface (MDI). Version 2.0.0 of the tool suite or higher was used for data analysis, contemporary with version 3.0.0 of the genomex-mdi-tools suite dependency and versions 1.3.2 and 1.8.2 of the mdi-pipeline-frameworks and mdi-apps-framework, respectively. Additional program dependency versions are set by the conda environment definitions tied to tool suite versions.

The following steps match previous pipeline descriptions ^35^: (i) read trimming, merging, and quality filtering using fastp ^93^, (ii) read alignment to the genome using bwa mem ^94^, (iii) aggregation of reads into read groups representing unique source DNA molecules, and (iv) SV junction detection using discordant alignment of paired reads and split reads. We applied the

‘align‘, ‘collatè, and ‘extract‘ pipeline actions to individual samples to discover potential discordant read alignments. The ‘find‘ action was then applied at once to all samples sequenced together in an experimental batch to find candidate SV junctions unique to one sample. The GRCh38/hg38 genome was used for all analyses and capture target regions were padded on each side by 800kb to set the allowable SV breakpoint regions.

Additions to the svCapture pipeline for this work relate to integrating results across multiple experimental batches, delivered by the ‘assemblè pipeline action and svCaptureAssembly Shiny app initialized in svx-mdi-tools v1.8.0. These tools apply standardized SV filtering, coverage assessments, and junction analysis, and assemble SV, target, and sample-level metadata into tables. We created individual assemblies for each cell line and another with all cell lines together. Most figures were generated using the svCaptureAssembly app working from those assemblies.

#### SV filtering

Filtering when counting SVs is essential to ensure that true, on-target SVs are counted in preference to read artifacts. Our goal throughout was to count only *de novo* SVs that arose during the experimental window prior to their expansion by replication. Accordingly, we only counted SV junctions found in a single source DNA molecule in a single sample as defined by the molecule outer endpoints. Those source DNA molecules needed to be sequenced by at least three read pairs to support their validity, since chimeric PCR artifacts arising late in PCR typically have only one matching read pair ^35^. Additional filters included (i) a required mapping quality of 30 or higher in one flanking alignment and 20 or higher in both, (ii) a requirement that at least one SV breakpoint fell in a capture target, with the other falling in a padded target region as defined above, and (iii) exclusion of deletion and duplication SVs less than 10kb or greater than 1.2Mb to match the established properties of SVs arising at CFSs ^17^. Inversions were filtered to exclude events less than 50kb due to a known artifact class in transposase libraries of false small inversions with large microhomology tracts resulting from intramolecular hybridization and synthesis during end-filling of Tn5-cleaved DNA ends (**Figure S1C**) ^35, 95^.

#### Target coverage assessment

Calculating effective target region coverage is essential for comparing samples. Unlike single-nucleotide variants (SNVs), not all bases in reads are equally able to report on the existence of a true SV. SV junctions cannot be detected near the ends of reads because a minimal span of approximately 20 bases must be properly aligned to the genome on each side of the junction.

Accordingly, we adjusted non-SV source DNA molecules to ensure that only bases that could have reported an SV junction were included in coverage calculations. Source molecules with fewer than three read pairs were excluded, and the length of the remaining molecules was adjusted by subtracting 2 x 20bp = 40bp from the actual length to disregard terminal bases where SV junctions could not be called. Target region coverage was calculated as the sum of all adjusted, on-target source molecule lengths divided by the summed length of all unpadded target regions. For visualization, base-level coverage was averaged over 100bp bins. Coverage is not uniform throughout target regions due to variable capture probe efficiency, read mappability, and underlying clonal SVs in cell lines. However, these systematic variations applied similarly to all samples of a cell line processed with the same capture probes.

### SV junction analysis

#### SV junction types and local structures

The svCapture pipeline reports without preference each of the four canonical types of SV junctions – deletions, duplications, inversions. and translocations ^1^ – where, as applied here, the first three types arise in a single capture target while translocations join two different targets.

Importantly, short reads can only detect junction sequences consistent with end joining; junctions arising through long blocks of homology are invisible to svCapture ^1, 35^. To characterize junctions, the last aligned bases nearest the junction on either side were defined as the two breakpoint positions in two coordinate systems: the reference genome and the SV-containing source DNA molecule. The distance between the junction positions in the reference genome defined the SV size. The junction was labeled as a microhomology event if the two junction positions overlapped in the source molecule such that the same read bases aligned to both reference breakpoints. The breakpoint positions of blunt joints abutted in the source molecule, whereas *de novo* insertions were evident as read bases between the breakpoint positions that did not align to either reference breakpoint. We plot insertions as positive separations of breakpoint positions in the SV molecule and microhomologies as negative separations, *i.e.*, overlapping alignments.

To compare junction profiles between experimental groups, we first calculated the average microhomology length of all junctions that did not have a *de novo* insertion, *e.g.*, R expression ‘mean(microhomologyLength[microhomologyLength >= 0])‘, to reveal the strand annealing preferences of the underlying mechanism(s). We further calculated the fraction of all junctions that had a *de novo* insertion between 2 and 15 bp, inclusive, to reveal the extent to which those mechanisms supported potentially templated insertions.

#### Insertion template discovery and characterization

Some novel bases inserted at junctions are known to be copied from template bases near the reference genome breakpoints ^45, 46, 65^. Locating templates is complicated by the fact that inserted sequences are often too short to be uniquely identified. However, base constraints when comparing a junction to a candidate template extend beyond the inserted bases to include “priming” and “resolving” microhomologies used during repair ^45^. To maximize sensitivity and specificity while searching for possible insertion templates, we adjusted the required number of flanking microhomology bases to ensure that a total of at least seven bases with one base of flanking homology on each side were used as a template query sequence. Thus, a single-base insertion was queried using three bases of initial microhomology on each side, a four-base insertion was queried using two bases of microhomology on each side, and insertions of five or more bases were queried with one base of microhomology on each side.

We searched for exact matches of the minimal search sequence for a given insertion junction in both strands of both reference genome breakpoints in a span 500bp upstream and downstream of the junction position. Thus, we searched widely over the reference bases that were both retained and lost at the junction over a total of 2 breakpoints x 2 strands x 2 junction sides x 500bp per side = 4kb genomic sequence. When a template match was found, the flanking microhomologies were expanded from the initial query to the maximum possible extent, stopping just before the first mismatch between the junction and template sequences. If multiple possible matches were found, we preferred the template with the longest span, including the flanking microhomologies, or, if that did not differentiate them, the template closest to the junction.

The type of the final selected insertion template, if any, was defined by its strand and placement in the reference breakpoints. As shown in figures, relative to a deletion SV junction, foldback and palindrome insertion templates were found in retained genomic sequence on the bottom genome strand, cross-junction templates were found in retained genomic sequence on the top strand, strand-switching templates were found on the bottom strand at least partially in the lost sequence, and slippage templates were found on the top genome strand in a manner that crossed the breakpoint junction position from retained into lost sequence. Importantly, retained and lost segments are defined relative to breakpoint positions, as defined above, not to the position of the underlying DSB ends, which cannot be ascertained from junction sequences.

### Quantification and statistical analysis

#### Number and source of experimental replicates

Throughout, figures are labeled to indicate the number of SVs or other relevant input counts that contributed to each plot. The number of samples contributing to each experimental group is evident from the plotted sample data points. Because of the time and expense involved in svCapture experiments, it was not possible to repeat all controls in all experimental batches. Accordingly, we plot all relevant data points that match an experimental group regardless of when they were acquired and indicate the shared experimental batches of different data points by their sample point color.

In general, our approach was to analyze sufficient replicates until the relationship between key experimental groups was established by statistical methods below. However, for completeness in reporting results, we sometimes show supplemental experiments performed in a single replicate with a given cell line. In such cases the data relationships match results from other cell lines and do not form the primary basis of data interpretation; single-replicate experiments should always be interpreted with caution and integrated in this way. Error bars on SV frequency plots represent the mean +/- 2 standard deviations or two or more data points.

#### Comparing intergroup SV Frequency

svCapture is a Poisson process in which an integer number of SVs of a particular class, most notably *de novo* deletions, are detected over an interval manifest as the library read depth, *i.e.*, more SVs are detected the more deeply a sample is sequenced. Thus, the Poisson rate for an svCapture sample is the number of detected SVs divided by the on-target coverage, noting that target regions were the same size in all samples from a given cell line. We label this Poisson rate “SV Frequency” to emphasize the intuitive sense that it approximates the fraction of aggregated target alleles in cells, *i.e.*, target haploid equivalents, that carried a *de novo* SV. Consistent with prior microarrays results ^17^ this fraction can exceed 50% of alleles under replication stress, reflecting the high mutational potential of the CFS hotspot genes we targeted.

Inter-sample differences in factors such as APH preparation potency, library insert sizes, fold target enrichment, and complexity are expected to influence SV count variance beyond random sampling, so we modeled svCapture data as an overdispersed Poisson using the negative binomial distribution (NBD). Moreover, svCapture results aggregate libraries prepared and sequenced over several years. Although we observed a high degree of reproducibility, batch effects such as unidentified differences in conditions and kits over time might also contribute to inter-sample variance.

To appropriately compare SV Frequency between experimental groups, *e.g.*, wild type vs. mutant, we performed pairwise intergroup comparisons using a generalized linear model based on the NBD in which the number of detected SVs, nSVs, varied as a function of the experimental group and batch as independent covariates. Target region coverage per sample was included as an offset parameter of slope 1 to effectively model SV Frequency. The relevant R language expression is ‘fit = MASS::glm.nb(nSvs ∼ group + batch + offset(log(coverage))‘, where ‘group‘ and ‘batch‘ are categorical variables and the p-value of the intergroup comparison was obtained from the ‘group‘ variable as ‘coef(summary(fit))[2, 4]‘. When glm.nb failed to return a result, the p-value was calculated using the glm function poisson model without overdispersion; such cases are denoted with a dashed significance line in plots. Throughout, we used p <= 0.01 as a significance threshold, with plot labels *, p <= 0.01; **, p <= 0.001; and ***, p <= 0.0001.

#### Assessing insertion template enrichment

Some insertions templates are expected to be found locally by chance at a Poisson rate of mu = 4kb search space x 1 / 4^nTemplateBases^. The probability of finding at least one local match corresponds to R expression ‘trialSuccessProb = 1 - dpois(0, mu)‘. To determine statistical significance of the actual number of found templates, we took each junction sequence at a given insertion size as an independent Bernoulli trial and estimated the p-value using R expression ‘1 - pbinom(nFound - 1, nSearched, trialSuccessProb)‘, where nSearched and nFound are the number of searched and found insertion templates, respectively, and the resulting p-value is the likelihood of finding nFound or more templates by random chance. A p-value <0.05 was taken as significant evidence for the contribution of local templates to the appearance of insertion junctions.

#### Cytogenetics data analysis

Data from experiments to examine the effects of MiDAS inhibition on APH-induced chromosome gaps and breaks and common fragile site expression were analyzed with the Student’s *t*-test.

## Data Availability

svCapture sequencing data have been deposited in two repositories consistent with human data access restrictions. Samples are listed in **Table S1**. Data from cell line UM-HF1 were deposited into the Database of Genotypes and Phenotypes (dbGaP, accession phs003121.v2.p1). Data from commercially available cell lines HCT116 and GM12878 were deposited into the Sequence Read Archive (SRA, BioProject ID PRJNA1085257). Microscopy data will be shared upon request.

Flow cytometry data were deposited into Mendeley Data (DOI 10.17632/mz53d2486n.1). The main processed data outputs of the svCapture pipeline are in **Tables S2 and S3**, included in the Zenodo code set linked to GitHub alongside the job scripts that generated them (see below), or in a separate Zenodo dataset carrying larger output files (DOI 10.5281/zenodo.10916987), including data packages and app bookmarks.

## Code Availability

Code comprising the svCapture data analysis pipeline and app is maintained in the svx-mdi-tools GitHub repository with all releases captured in Zenodo (DOI 10.5281/zenodo.10928148). Data-specific job scripts used to execute the pipeline for samples in this manuscript and associated support files, including resource files, sample lists, and job logs, are maintained in a separate GitHub repository linked to Zenodo (DOI 10.5281/zenodo.10936173). Together, data and code above can reproduce our analyses.

## Acknowledgements

This work was funded by grants CA200731 and GM147026 from the National Institutes of Health to T.E.W. and T.W.G. We thank Dale Ramsden for reagents, advice in establishing TMEJ/NHEJ assays, and for valuable discussion. We thank Pamela Bennett-Baker for assistance with reagent preparation and method validation in early stages of the work and Charles Kazazian for assistance with cytogenetics. We thank the University of Michigan Advanced Genomics Core for skilled handling of svCapture library preparation and sequencing, and the University of Michigan Flow Cytometry Core for expert assistance with long flow sorting sessions. We thank Patrick O’Brien and Martin Arlt for critical reading of the manuscript and valuable input over many years.

## Author Contributions

**T.**E.W. designed the experiments, wrote the data analysis pipeline, analyzed data, and wrote the manuscript; S.A. designed and performed the experiments and analyzed data; A.W. assisted in the design and performance of experiments; T.W.G designed the experiments, performed cytogenetic analysis, analyzed data, and wrote the manuscript.

## Materials & Correspondence

Further information and requests related to cell resources and wet lab reagents should be directed to Thomas W. Glover. Requests related to data analysis and bioinformatics tools should be directed to Thomas E. Wilson.

## Supplemental Tables

**Table S1. svCapture samples.**

svCapture samples with project batches, samples names and identifiers, and experimental conditions.

**Table S2. svCapture junction properties.**

Properties of individual SV junctions of all types from all samples.

**Table S3. svCapture sample group summary.**

Aggregated experimental results for all groups of replicate samples, deletion SVs only.

**Table S4. CRISPR/Cas9 guides and alleles and TMEJ-NHEJ assay.**

Excel file with three spreadsheets listing details of (i) CRISPR guide RNAs, (ii) CRISPR KO alleles, and (iii) oligonucleotides used in the TMEJ-NHEJ assay.

