## Supplemental Information for "POLQ mediates replication-stress induced structural variant formation throughout common fragile sites during mitosis"

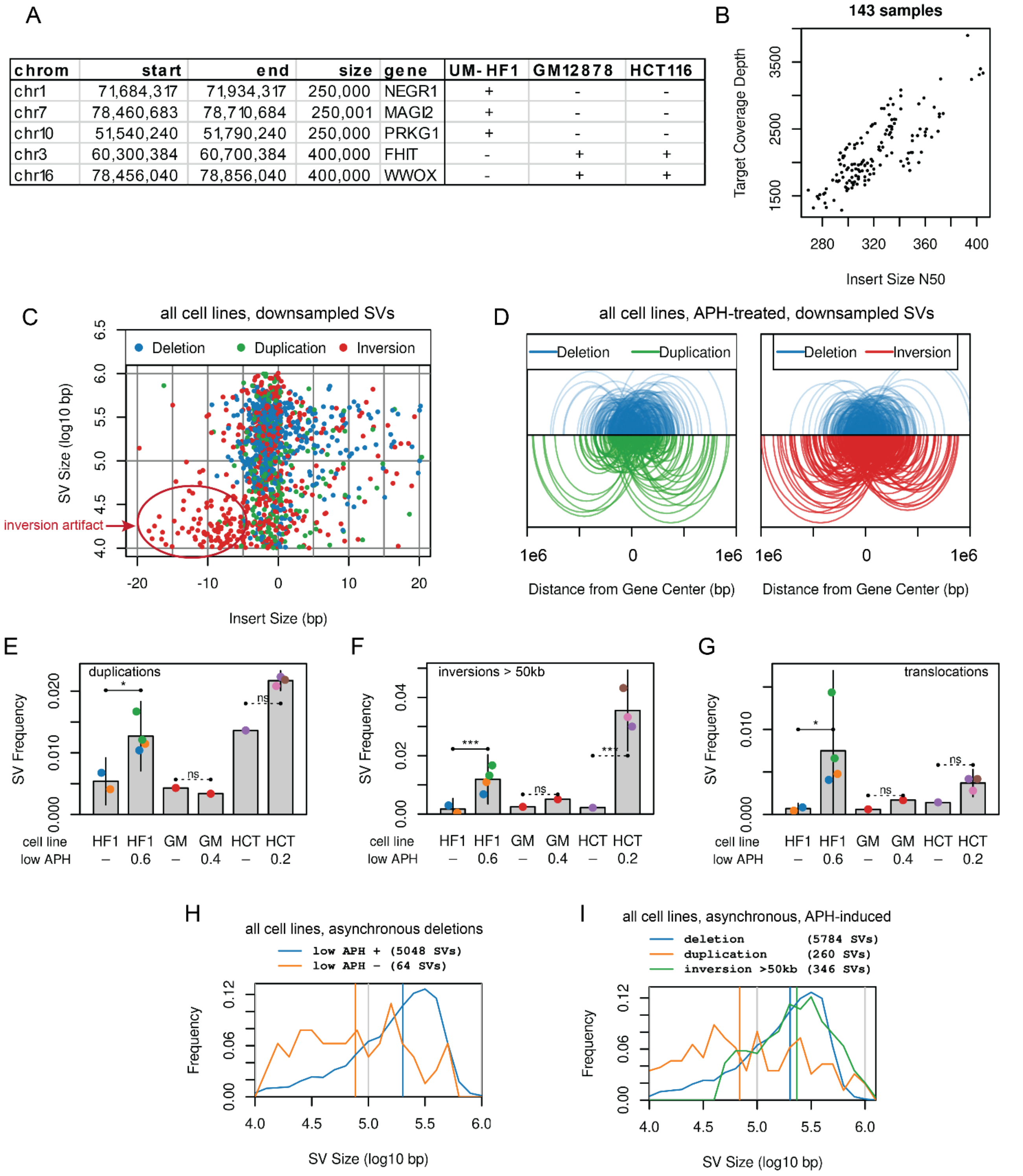


| chrom | start | end | size | gene | UM-HF1 | GM12878 | HCT116 |
|---|---|---|---|---|---|---|---|
| chr1 | 71,684,317 | 71,934,317 | 250,000 | NEGR1 | + | - | - |
| chr7 | 78,460,683 | 78,710,684 | 250,001 | MAGI2 | + | - | - |
| chr10 | 51,540,240 | 51,790,240 | 250,000 | PRKG1 | + | - | - |
| chr3 | 60,300,384 | 60,700,384 | 400,000 | FHIT | - | + | + |
| chr16 | 78,456,040 | 78,856,040 | 400,000 | WWOX | - | + | + |

**Figure S1. svCapture profiles in asynchronous human cell lines.**

**A.** svCapture target regions used in different cell lines (“+” symbols).

**B.** Summary of library insert sizes, expressed as N50, and target region coverage over all samples.

**C.** Correlation of SV size to junction insert/microhomology size by SV type. Each type is downsampled to the inversion event count for comparison. Circled small inversions with large junction microhomologies are a known library artifact resulting from intramolecular hybridization and synthesis during end-filling of Tn5-cleaved ends. Accordingly, only inversions >50kb are included in subsequent figures.

**D.** Arcs connect the two breakpoint positions of 500 individual SVs of each type. Color intensity reflects the fraction of SVs at that size to emphasize that deletions arise more central to CFS genes whereas duplications and inversions have a higher fraction of flanking events.

**E to G.** Duplication, inversion, and inter-target translocation SV induction, respectively. Note the Y-axis limits and lower fold induction as compared to deletions in Figure 1.

**H.** SV size distributions merging asynchronous cultures of all cell lines, stratified by low-dose APH treatment to show the larger size of induced deletions.

**I.** SV size distributions from asynchronous, APH-treated cells, now stratified by SV type to show the smaller size of duplications. With lower fold induction, more duplications are expected to have arisen prior to APH treatment.

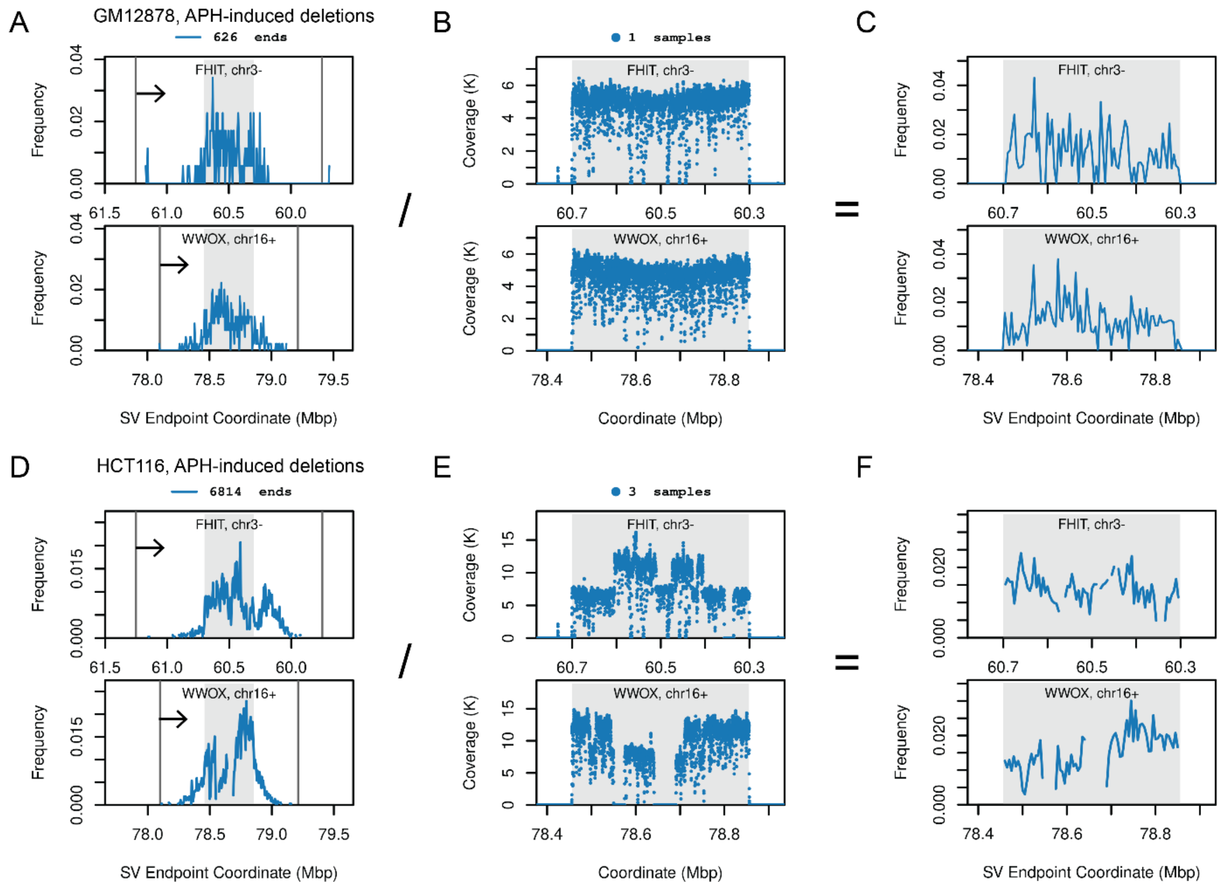


**Figure S2. SV breakpoint locations in asynchronous human cell lines.**

**A to C.** Placement of GM12878 APH-induced deletion breakpoints (A and C) and read coverage (B) in the *FHIT* and *WWOX* target genes.

**D to F.** Like panels A to C, now for HCT116 cells. Panels C and F show breakpoint distributions from panels A and D normalized to read coverage >= 500 from panels B and E. Panel E reveals clonal baseline SVs in HCT116 at *FHIT* and *WWOX*, consistent with prior genomic analysis of this cell line. Additional *de novo* SVs still occur superimposed onto the alleles carrying these baseline SVs.

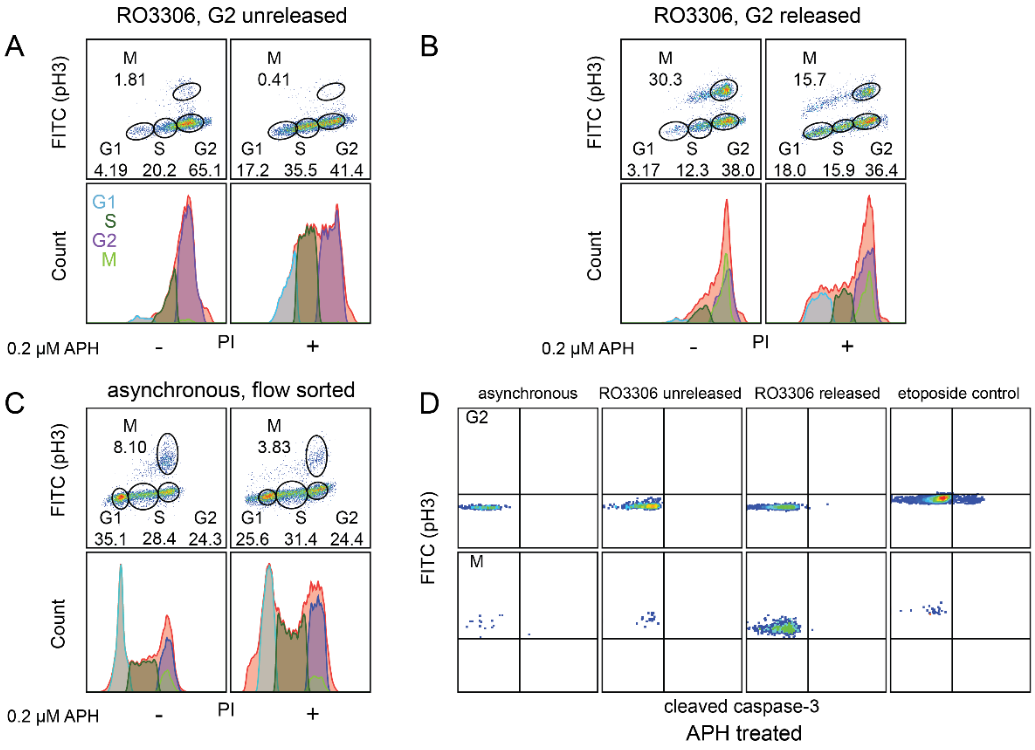


**Figure S3. Isolation and viability of S, G2, and M-phase cells.**

**A.** Example HCT116 flow sort for the synchronization paradigm with G2 cells collected prior to release from RO3306. x-axis, DNA content; y-axis, pH3. Lower panels show cell cycle distributions, revealing synchronization and the impact of APH on S-phase accumulation.

**B.** Like A, now for the RO3306 synchronization paradigm with G2 cells collected after release.

**C.** Like A, now for HCT116 cells flow sorted by cell cycle phase from cultures without RO3306.

**D.** Flow cytometry plots of cleaved caspase 3, x-axis, and pH, y-axis, showing that G2 cells remained viable during APH treatment, arrest, and release. The rightmost panel was treated with 10µM etoposide as a positive control.

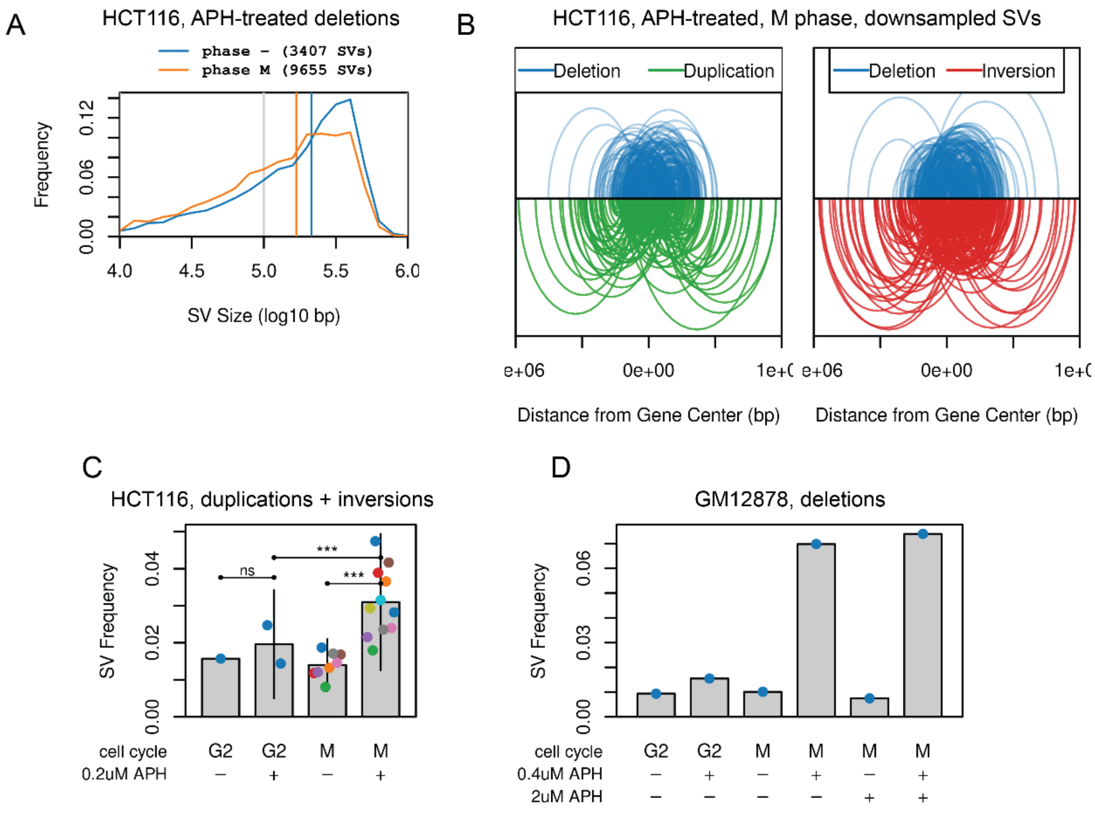


**Figure S4. APH induces SV junction formation primarily during mitosis.**

**A.** Similar SV length distributions for APH-induced deletions in asynchronous vs. M-phase HCT116 cells.

**B.** Like Figure S1D, showing that deletions induced in M-phase HCT116 cells tend to be more central as compared to duplications and inversions.

**C.** Duplication plus inversion frequencies as a function of cell cycle phase in HCT116.

**D.** SV frequencies from GM12878 cells harvested with and without high-dose APH suppression of MiDAS. This single replicate experiment matches results from HCT116.

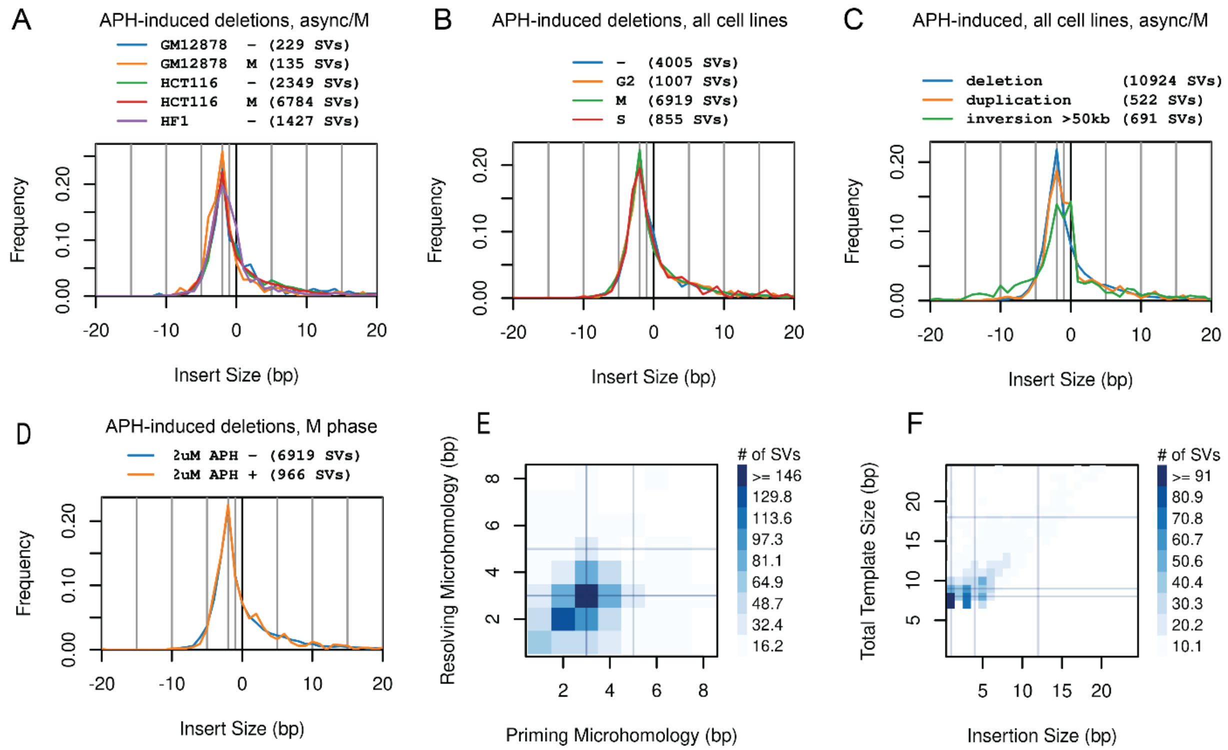


**Figure S5. Junction analysis for microhomologies and templated insertions.**

**A.** Distribution of microhomology and insert sizes by cell line and cell cycle phase (async/- vs. M).

**B.** Like panel A, now by cell cycle phase over APH-treated samples from all cell lines.

**C.** Like panel A, now by SV type detected in asynchronous or M-phase cells for all APH-treated samples.

**D.** Like panel A, now showing APH-induced deletions in M-phase by concurrent high-dose APH treatment. Distributions throughout are nearly identical, except for inversions that likely retain a small amount of known artifact mechanisms unique to that SV type.

**E.** Correlated size distributions of priming and resolving microhomologies over all insertion templates.

**F.** Correlated insert and total template (including flanking microhomologies) size distributions.

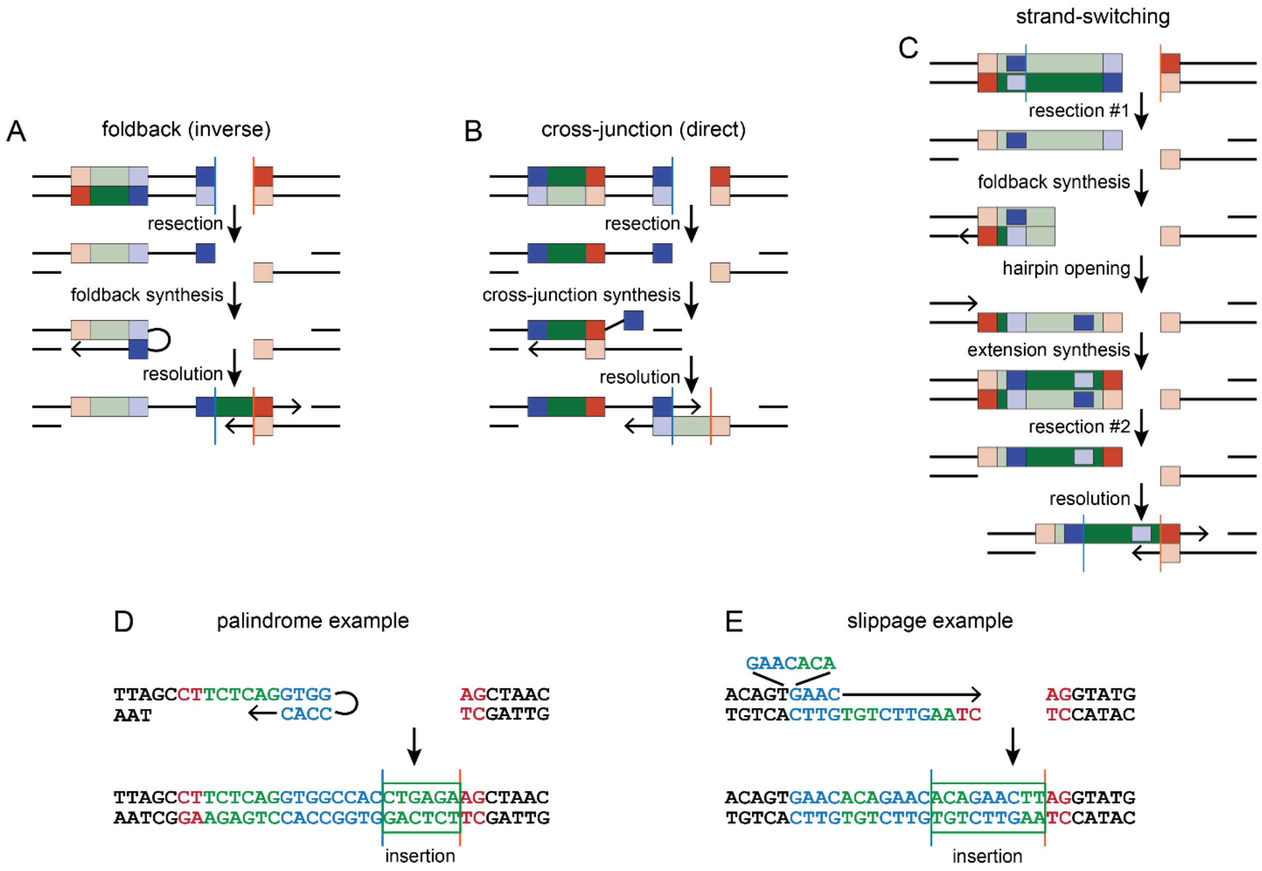


**Figure S6. Mechanisms of templated insertion.**

**A** to **C.** Previously described templated insertion mechanisms. Only left-sided templates are drawn; right-sided templates are flipped but retain the same strandedness, *e.g.*, right-sided foldback templates are also on the bottom strand. The strand-switching mechanism was not readily observed in our data.

**D.** Example of a small subset of foldback events where the inferred hairpin structure was a perfect palindrome, which alters the placement of the called junction position as compared to non-palindromic foldbacks. This pattern is most likely a coincidence rather than a distinct insertion mechanism.

**E.** Example of an observed slippage-class insertion. Base colors and boxes in D and E follow Figure 4A.

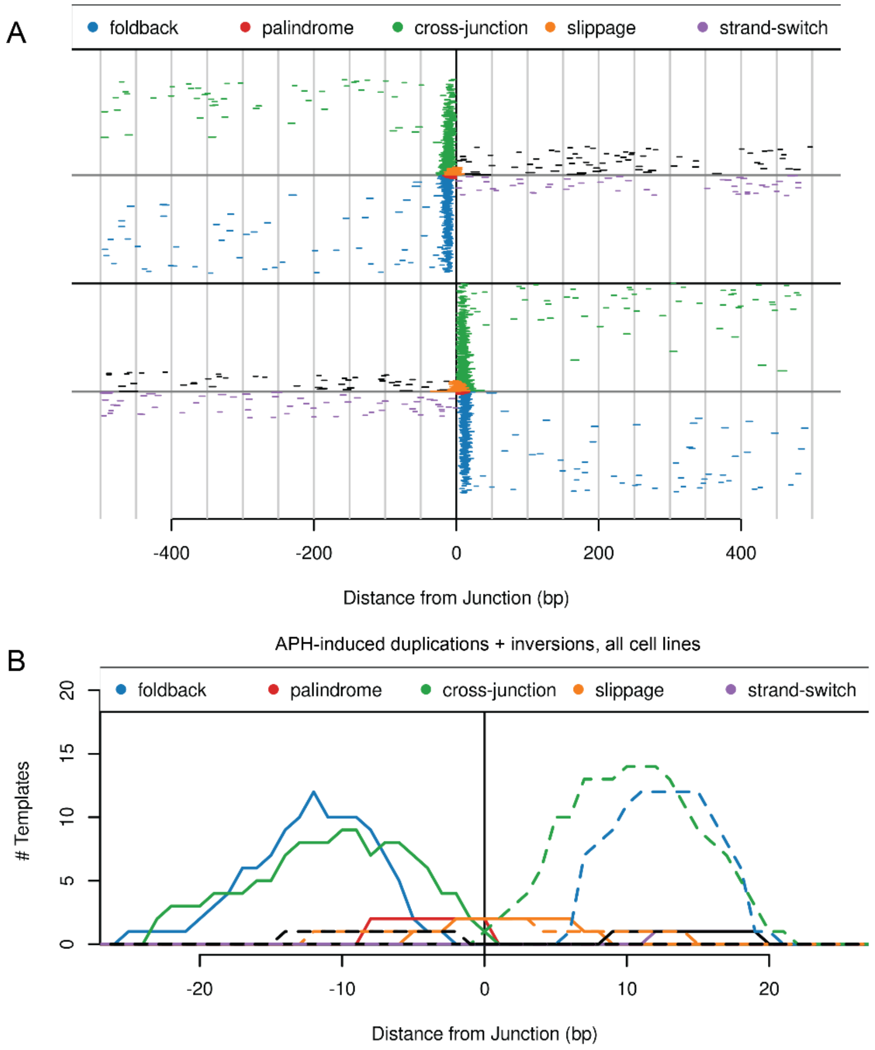


**Figure S7. Locations of insertion templates.**

**A.** Wide view of insertion template locations for APH-induced deletions throughout the 1kb interrogation regions. Randomly distributed candidate templates further from breakpoints likely represent fortuitous matches unrelated to junction formation mechanisms.

**B.** Bases contributing to insertion templates at APH-induced duplication and inversion SVs across all cell lines, showing a similar template location pattern as for deletions.

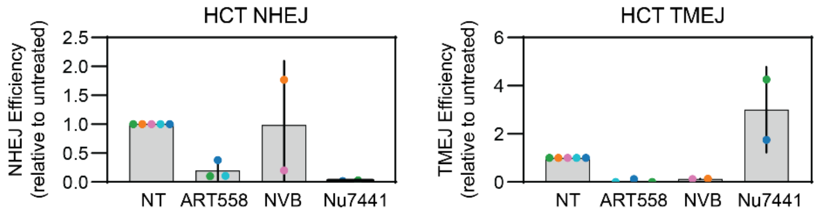


**Figure S8. Validation of DSB repair inhibition.**

Transfected oligonucleotide assay assessing inhibition of NHEJ (left) and TMEJ (right) by DNA repair inhibitors.

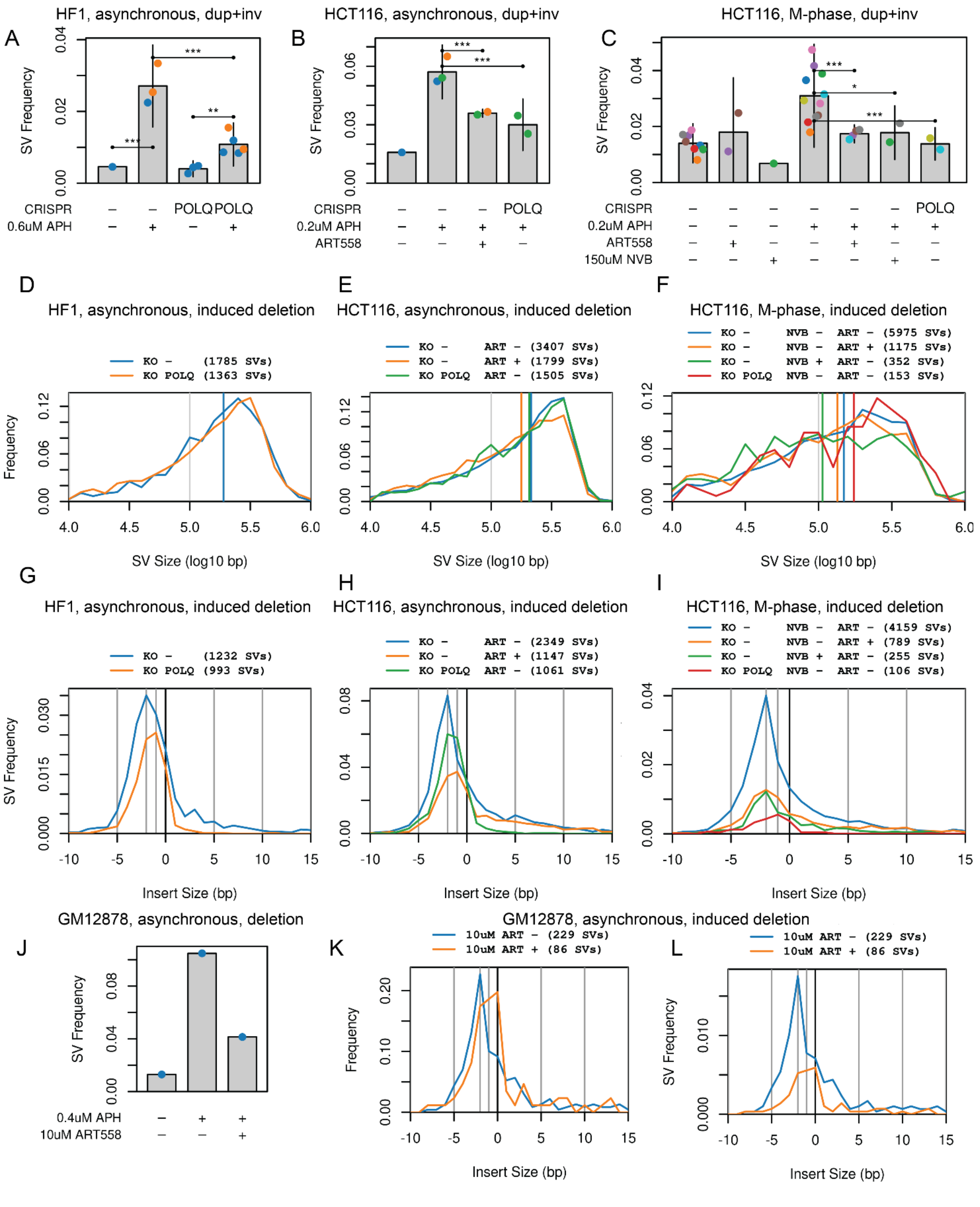


**Figure S9. POLQ role in SV formation at CFSs.**

**A** to **C.** SV frequency plots showing duplication plus inversion SVs.

**D** to **F.** Induced deletion SV size distributions for the data in Figures 5D to 5F, which are not altered by TMEJ suppression.

**G** to **I.** Insertion/microhomology sizes of the same data as Figures 5G to 5I, now normalized to the summed target region coverage in each group. Thus, the y-axis scale of these plots is the same as the SV Frequency plots to reveal the absolute formation efficiency of different SV junction types.

**J.** Partial inhibition of deletion SV formation in asynchronous GM12878 cells by ART558. This single replicate experiment matches results from HF1 and HCT116 cells.

**K.** Insertion/microhomology size distributions for the data in panel J.

**L.** The same data as panel K normalized to the summed target region coverage as in panels G to I.

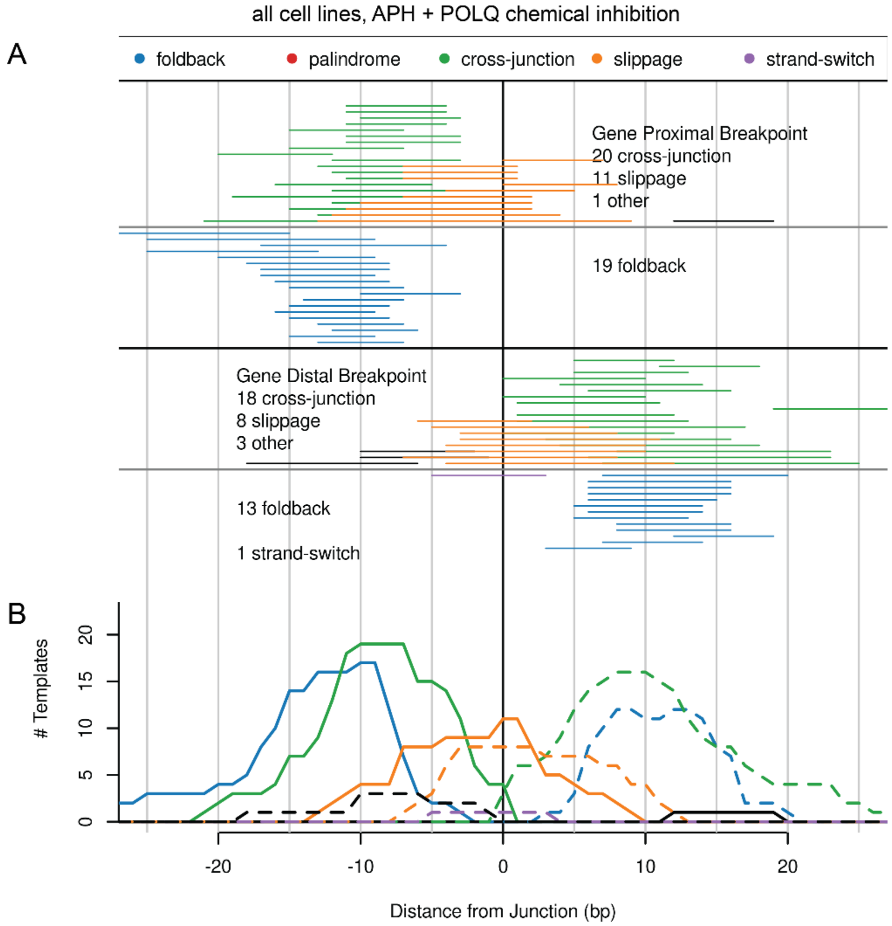


**Figure S10. Insertion template locations under POLQ inhibition.**

**A** and **B.** Pileup and histogram plots like Figures 4F and 4G, respectively, showing insertion template locations for all cell lines treated with low-dose APH plus either NVB or ART558 to inhibit POLQ. A larger fraction of templates are of the slippage class under POLQ inhibition, but cross-junction and foldback templates are observed.

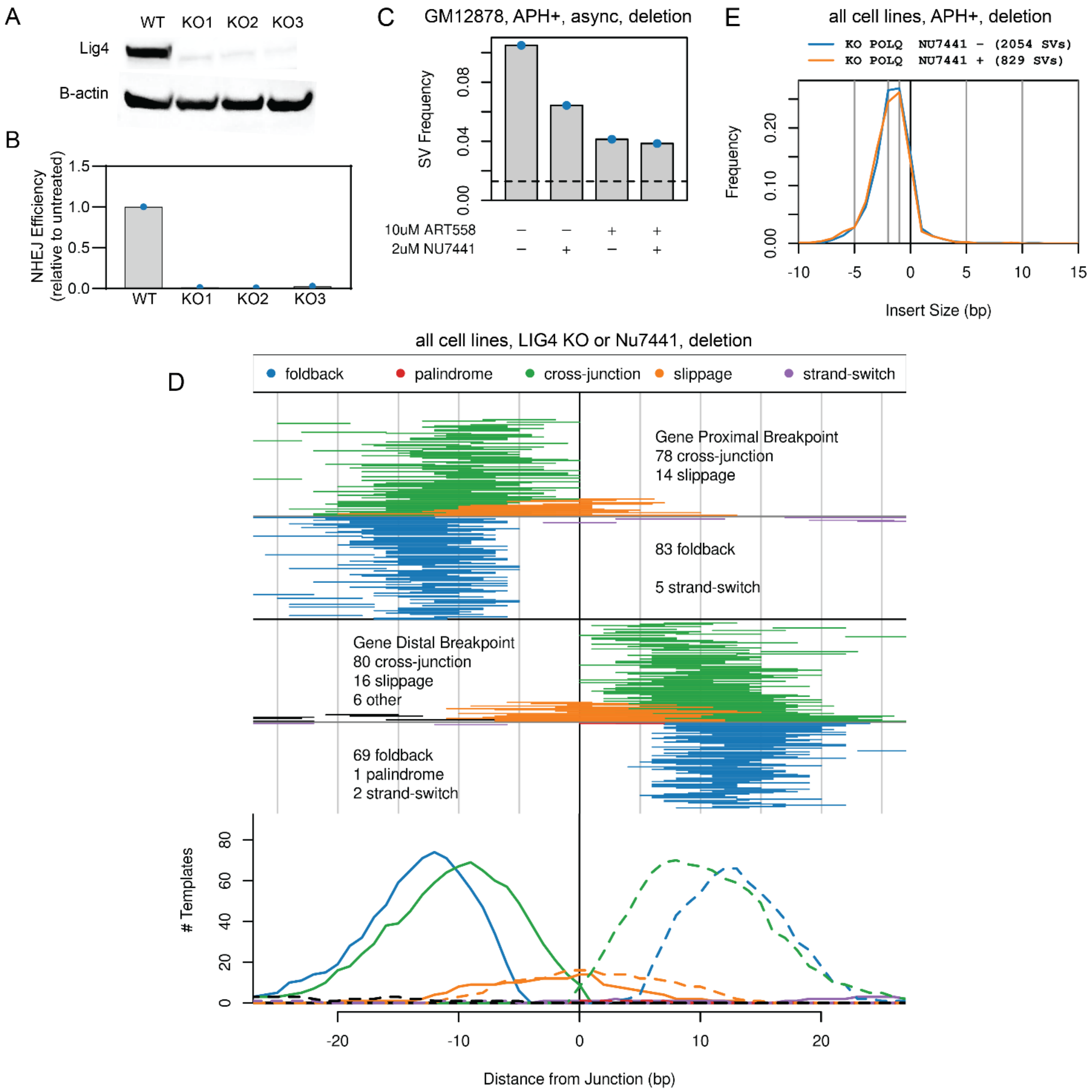


**Figure S11. Relationship between TMEJ and NHEJ in SV formation.**

**A** and **B.** Western blot and NHEJ assay with transfected oligonucleotides, respectively, verifying *LIG4* KO in HCT116 cells.

**C.** Effect of TMEJ and NHEJ chemical inhibition on deletion formation in APH-treated GM12878 cells.

**D.** Plots showing unaltered insertion template locations in samples lacking NHEJ due to either *LIG4* knockout or treatment with NU7441 as compared to data in Figures 4F and 4G.

**E.** NHEJ loss does not alter junction microhomology and insertion profiles in the absence of POLQ.
